# The Role of Attention in Multi Attribute Decision Making

**DOI:** 10.1101/2023.12.06.570439

**Authors:** Aaron Sampson, You-Ping Yang, Marius Usher, Dino Levy, Ernst Niebur, Veit Stuphorn

## Abstract

Real-life decisions typically involve multiple options, each with multiple attributes affecting value. In such complex cases, sequential shifts of attention to specific options and attributes are thought to guide the decision process. We designed a task that allowed us to monitor attention in monkeys engaged in such multi-attribute decisions. We recorded pre-supplementary motor area neurons encoding action value signals reflecting the decision process. Attention guides this process through two mechanisms. First, attention enhances the activity of neurons representing the currently sampled option, independent of the attended option value. Second, attention up-regulates the gain of information integration towards the evolving value estimate for the attended option. In contrast, we found no evidence for a third suggested mechanism, in which only the attended option is represented. Instead, attention influences the ongoing information accumulation and competition between the options by modulating the strength of the value information that drives this circuit.

## 1 Introduction

In many real-world decisions, humans and animals have to choose between multiple options whose value is influenced by multiple attributes. For example, humans might have to choose between different cars that vary in maker, price and fuel efficiency. Likewise, non-human primates searching for food might have to decide between different items that vary in nutrient content, availability, and required effort [1]. An optimal decision strategy requires integration of information about each attribute independently for each option, before the values of the options are compared. However, behavioral experiments in humans and animals indicate that option values are not estimated independently, and that choices often deviate systematically from optimality [2, 3].

It is thought that this is because an optimal decision strategy would require the representation and integration of large amounts of information. To overcome this cognitive resource limitation, the attentional system is thought to select only some of the available information that is then used to dynamically construct preferences. Indeed, decisions in real-world situations are typically preceded by multiple shifts of attention between available options [4]. Behavioral experiments have shown correlations between attention and choice [5], and neuronal recording experiments have started to investigate the role of attention in decision making [6–9]. A recent study has found strong effects of attention shifts on value and choice-related signals in frontal areas [6]. However, the exact mechanism by which attention influences decision making remains unclear.

At the neural level, three mechanisms have been proposed for how attention affects decision-making: the ‘additive’, ‘gain-modulation’, and ‘serial’ models (Fig. 1). The first two assume competing pools of neurons that reflect the current value estimate of their preferred option and are influenced by attention in two different ways (Fig. 1A). First, in the ‘additive’ model (yellow), attention to an option provides separate inputs into the different value representing pools, which add to the subjective value of the attended option and subtract from the value of all unattended ones [10, 11]. The model predicts that neuronal activity shows a binary shift in activity levels that reflects which option is attended, but not its value (Fig. 1B). Attention to an option thus increases the likelihood that it is chosen independent of any of its attributes.

**Fig. 1:**
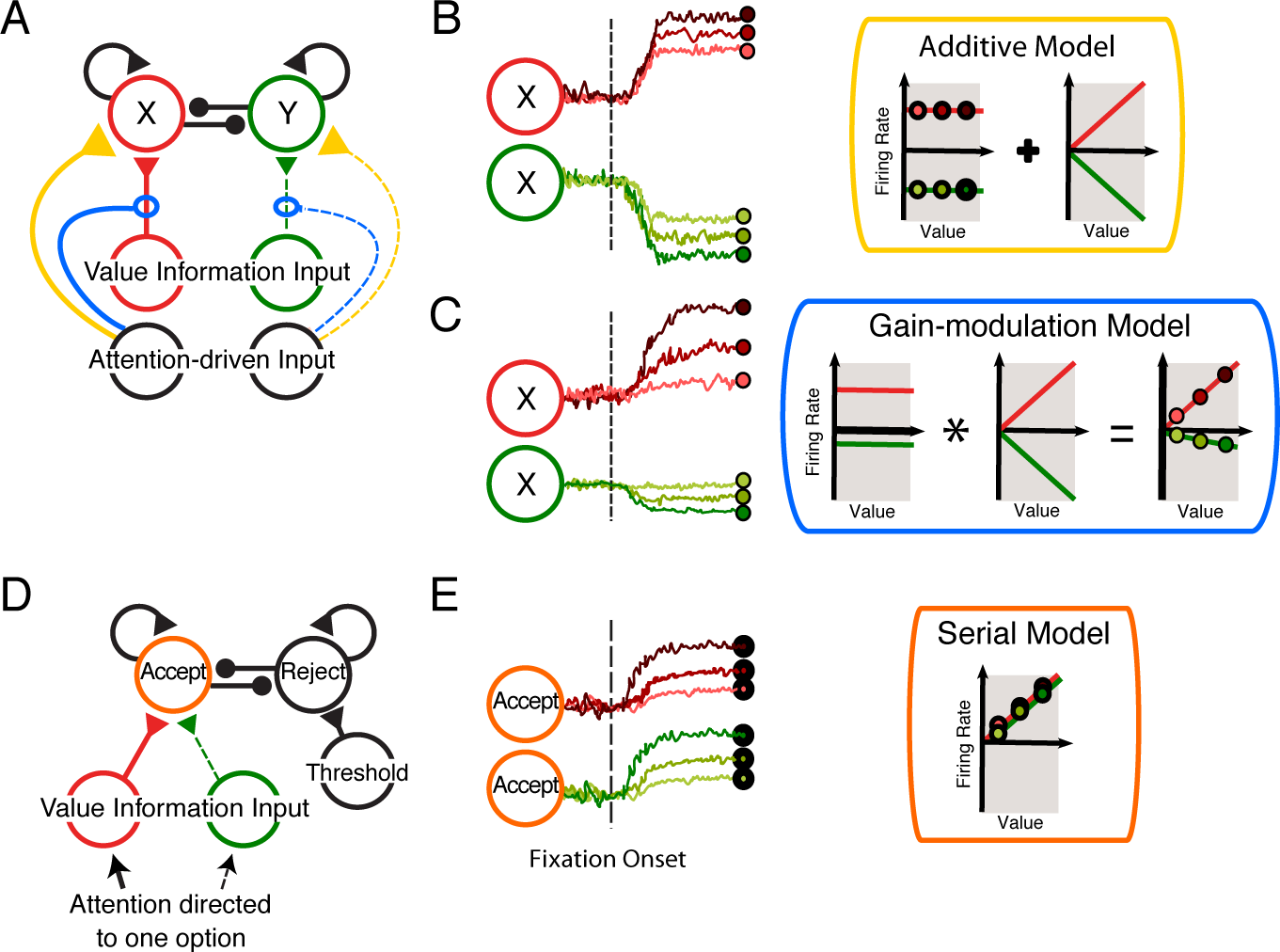
Three possible roles for attention in decision-making. (A) Standard decision architecture consisting of mutually inhibitory pools of neurons representing each of two alternatives (options X and Y), with their activity being driven by value-related inputs. Additional, attention-driven input modulates activity in the competing pools of value-encoding neurons. Two forms of modulation are possible: Attention can provide added input to value-encoding neurons (Additive Model; yellow connection) or change the gain of their input (Gain-modulation model; blue connection). In either case, whether an option is attended (solid line) or unattended (dashed line) has opposite effects. (B) Predicted effects of additive modulation on value signals. Left: Schematic activity of neurons encoding the value of option X as function of time when option X is attended (red) vs. when option Y is attended (green). Fixation onset is indicated by the dashed vertical line. Dark to light shading indicates high to low value of the attended option. Right, inside yellow frame: Schematic of model components. Attentional influence (left graph) is independent of option value (shading) and linearly combined with option value (right graph). Because attention is an independent factor that is added to option value its influence on neuronal activity can be linearly separated. (C) Predicted effects of gain-modulation on value signals, conventions as in (B). Because attention multiplicatively changes the gain of the response to value, its effect cannot be separated from value. Because gain depends on the type of attended option, the scale of the value response is predicted to be different for the two options. (D) Alternative decision architecture, in which mutually inhibitory pools of neurons represent the option to accept or to reject the currently attended option. The accept option is driven by the value of the currently attended option. The reject option is driven by a threshold for the minimal value level that is acceptable. Attention is directed serially to the available options and decisions are a series of accept/reject choices. (E) Predicted effects of a serial decision architecture on value signals. Because attention controls representation, only the value of the currently attended option is represented.

Second, in the ‘gain-modulation’ model (blue), the importance of the attended evidence for estimating an option’s value is increased. This in turn increases the gain of the value representation for the attended option and decreases it for the unattended one [12]. Attention to an option does not always increase the probability that it is chosen, because it can both increase or decrease its value depending on the attended information. Instead, attention influences decisions indirectly by changing the strength with which decision-relevant information affects preference formation. The model therefore predicts that neuronal activity reflects the value of attended and unattended options with different gains (Fig. 1C).

Third, in the serial model (orange), only the currently attended option is represented by the decision-making circuitry, even if multiple options are present concurrently [13, 14] (Fig. 1B). Hence, the attended option does not directly compete with alternative options. Instead, it competes with an acceptance threshold, and is either accepted or rejected. Rejection leads to a shift of attention to another option, and this procedure continues until an option is accepted. Thus, the model predicts that only the value of the currently attended option is represented, independent of its position (Fig. 1E).

We developed an attention-guided multi-attribute decision-making task that allows us to observe the shifting focus of attention of macaque monkeys while they evaluate the options and select one.We recorded single unit activity in medial frontal cortex, specifically the pre-supplementary motor area (preSMA). Neuronal activity in medial frontal cortex reflects motor readiness, action value, and action selection of eye and arm movements in humans and monkeys [15–18].

We confirmed that preSMA neurons represent the expected values of choices selected by arm movements to preferred directions, and we showed that these arm-movement specific action value signals change dynamically as the monkeys evaluate the attributes of the two options. This allowed us to use our knowledge of the current allocation of attention at each fixation to investigate the influence of attention on the ongoing decision process and to compare our results with predictions of the three discussed models. Our findings support the additive and gain-modulation models, but argue against the serial model.

## 2 Results

Two monkeys (*Macaca mulatta*, I and H) were trained to perform a multiattribute decision-making task (Fig. 2A) that required sequential sampling of options and attributes. The task clearly demarcates the inspection of each attribute and option, allowing us to identify the neural activity that corresponds to the sampling of each piece of decision-relevant information. The monkeys were free to make as many fixations and in whatever order as needed to gather sufficient information to make a decision. Because the task is self-terminating, it provides a behavioral indication of when sufficient information has been gathered to reach a decision.

**Fig. 2:**
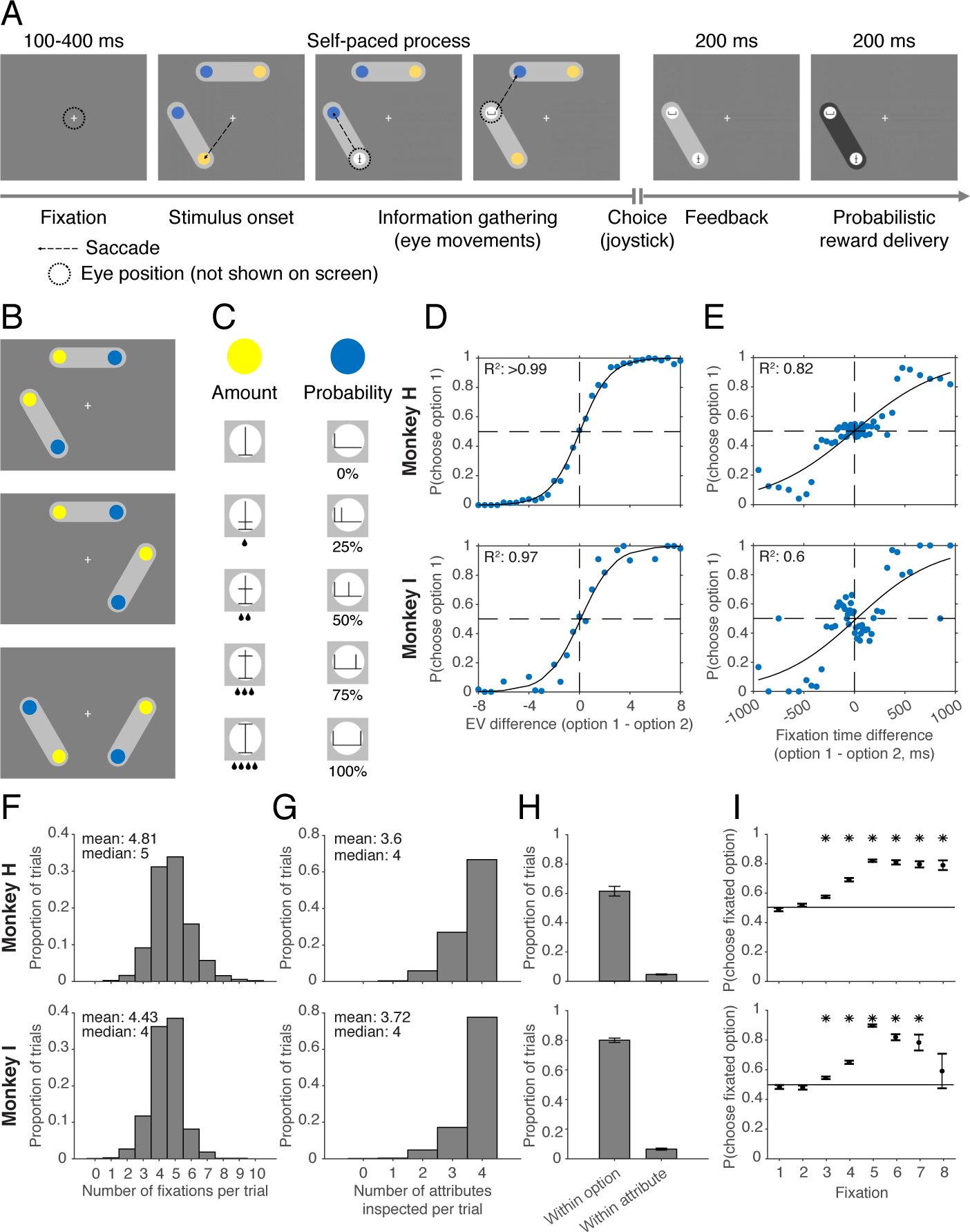
Multi-attribute decision-making task. (A) Task structure. After trial initiation, two options are represented by light gray ellipses within which colored circles (masks) indicate their two attributes: reward amount (yellow) and probability (blue). The monkeys are free to direct their gaze to explore the two options. When the monkeys fixate an attribute, the mask is removed, and the revealed symbol shows the attribute value. The monkey chooses by moving a joystick in the direction of the chosen option (here left). This option is then shown by itself after which the background color of the ellipse changes from light to dark gray and the outcome is delivered (depending on its reward amount and probability). (B) In each trial, the options were presented randomly in one of three possible spatial configurations. (C) Amount attributes could take values 0, 1, 2, 3, or 4 (corresponding to the volume of the water reward), and probability attributes could take on values 0, 0.25, 0.5, 0.75, or 1, corresponding to the likelihood of receiving a water reward. (D–I) Behavioral characteristics for Monkey H (top) and Monkey I (bottom). (D) Probability across all trials of choosing option 1 as a function of the difference in expected value between the two options. Blue points are empirical choice probabilities computed from behavioral data across recording sessions, black curve is a logistic fit. (E) Probability across all trials of choosing option 1 as a function of the difference in total time spent fixating the two options. Blue points are empirical choice probabilities computed from behavioral data across recording sessions, black curve is a logistic fit. (F) Number of fixations made per trial (Monkey H: 76,766 fixations from 15,971 trials; Monkey I: 25,935 fixations from 5,850 trials). (G) Number of unique attributes sampled per trial. (F,G) Red bars correspond to trials in which a zero-value option was presented (either amount or probability equal to zero). (H) Proportion of trials in which the first transition between attributes is within an option, or within an attribute (across options). (I) Probability of choosing the currently fixated option for sequential fixations within trials. Fixations with a mean choice probability across recording sessions significantly different from chance (one-sample t-test, *p <* 0.05) are denoted with an asterisk at the top of the axes. (H,I) Mean across recording sessions *±* SEM).

On each trial, the monkeys had to choose between two options with different combinations of two attributes: reward amount (*a*) and probability (*p*). The two options were displayed in two of three possible positions (Fig. 2B). The attributes of both options were masked behind colored circles, which indicated the type of attribute. The animal’s eye movements were tracked, and only when it fixated an attribute, the corresponding colored circle was removed and a cue indicating the attribute’s magnitude was presented (Fig. 2C). Animals indicated their decision by moving a joystick in the direction of the chosen option.

### 2.1 Information Sampling and Choice Behavior

The monkeys understood the meaning of the symbolic cues (see Suppl. Material). On most trials, the monkeys chose the option with higher expected value (EV) (Fig. 2D), indicating that the monkeys integrated and compared the attributes for both options. In human and monkey free-viewing tasks, choice and the amount of time inspecting an option are correlated [19–21]. We found a similar tendency of the monkeys to fixate the attributes of their chosen option more than those of the other option. This correlation allowed the monkeys’ choice to be predicted purely based on the difference in total fixation time between the two options (Fig. 2E), albeit a weaker prediction than that based on the difference in expected value.

Before a choice, the monkeys needed to sample the cues. To determine whether the monkeys used all available information, we examined how many fixations the monkeys made and how many cues among the four attributes they sampled before choosing (Fig. 2F,G). In the majority of trials, both monkeys made at least four fixations (#fix*≥*4: Monkey I: 85.11%; Monkey H:88.86%; Figure 2F) and sampled all available information (#attributes sampled=4: Monkey I: 77.59%; Monkey H: 66.75%; Figure 2G). They had a very strong preference for inspecting both attributes of one option (within-option sampling), before doing the same with the other option (Fig. 2H). Consequently, they showed no bias towards choosing the first inspected option and only started to commit after inspecting both options (Fig. 2I). Taken together, the monkeys’ choice and fixation behavior show that they typically based their decisions on the inspection of all attributes.

### 2.2 PreSMA Neurons Represent the Developing Decision Process

We recorded neural activity of 84 preSMA neurons (monkey H: 62; monkey I: 22) while monkeys performed the attention-guided multi-attribute decision task. Fig. 3A–B shows recording locations overlaid on MRI. The preSMA neurons encode Attention-, Value-, and Choice-Related Signals (Fig. 3C–D). The activity of many neurons in preSMA was correlated with the monkeys’ chosen arm movement and the value of the chosen option (Fig. 3C). We refer to the arm movement direction that evokes the strongest activity as the preferred direction (PD) of each neuron, the other two directions were defined as non-preferred directions (NPDs). Each of the three arm movements was significantly preferred by a substantial number of preSMA neurons (Up: 22/84, 26%; Left: 26/84, 31%; Right: 11/84, 13%; ANOVA, *p <* 0.05). These signals often appeared long (*>* 600 ms) before the onset of the arm movement, when the monkeys were still inspecting option attributes. The spatial selectivity of the motor response provides a framework for understanding the information represented by the neurons during the fixation of specific attributes (Fig. 3D). The activity of many preSMA neurons reflects whether the attended spatial position matches their preferred direction or not (Attention). In addition, preSMA neurons also carry signals about the value of the inspected options in these earlier time periods. Thus, preSMA neurons encode a premotor signal that indicates the evolving likelihood that a specific arm movement will be generated, given the information gained from a sequence of inspected attributes. While some neurons seem to only reflect one of the variables, most neurons clearly reflect more than one variable.

**Fig. 3:**
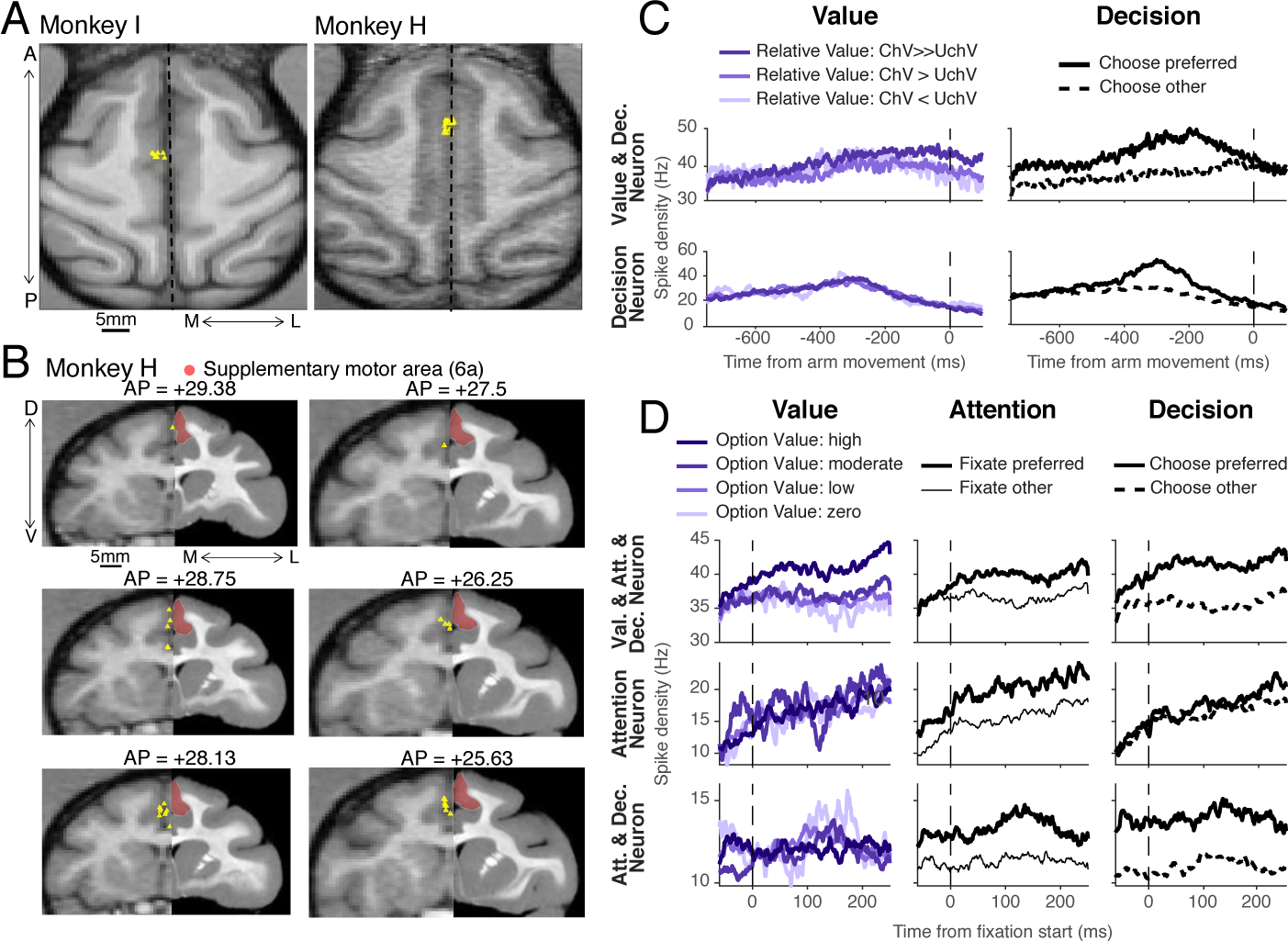
MRI reconstruction of SMA/pre-SMA neuron recording locations. (A) Top view of the recording sites of monkey I and monkey H, respectively. (B) Coronal view of recording sites in monkey H. Yellow triangles indicate recording locations, red area indicates area 6a. (C–D)Pre-SMA neurons encoding value, attention, and decision. Selected neurons are labelled according to the variable they seem to be representing most clearly. Each panel shows the instantaneous firing rate of one neuron (row), averaged over all traces defined by setting one variable (legend above top row) to a specific condition (column). (C) Neuronal activity aligned on arm movement onset. The first column shows activity sorted by the value of the chosen option (ChV) relative to the one of the unchosen option (UchV). The second column shows activity sorted by whether the chosen direction was in these neurons’ preferred direction (PD; ”Choose preferred”) or one of their non-preferred directions (NPDs; ”Choose other”). (D) Neuronal activity aligned on the start of fixation of attributes. The first column shows activity sorted by option value (light to dark for low to high value). The second column shows activity sorted by attention (thick lines: fixation of option in the PD, thin lines: fixation in an NPD). The third column shows activity sorted by decision (solid lines: chosen option in the PD, dashed lines: chosen option in an NPD).

We searched for preSMA neurons whose changing activity reflects the decision process. To identify preSMA neurons encoding the decision and its expected value, we used generalized linear models (GLMs) (Extended Data Table 1). On each trial two distinct pools of preSMA neurons represent the action values of the two current options. The action value signals consist of two components. First, the presence of *V_dp_* signals (65/84, 77%) indicates that the preSMA neuron pools robustly represent the value of their respective PD option, independent of the value of the other one. Second, the *RV_d_p* signal (61/84, 73%) indicates that the pools also partially compete with each other, because increased value of the NPD option reduces action value signals for the PD option. This suggests a parallel, comparative decision mechanism similar to what has been described in other cortical areas [15, 16, 18, 22].

**Table 1:**
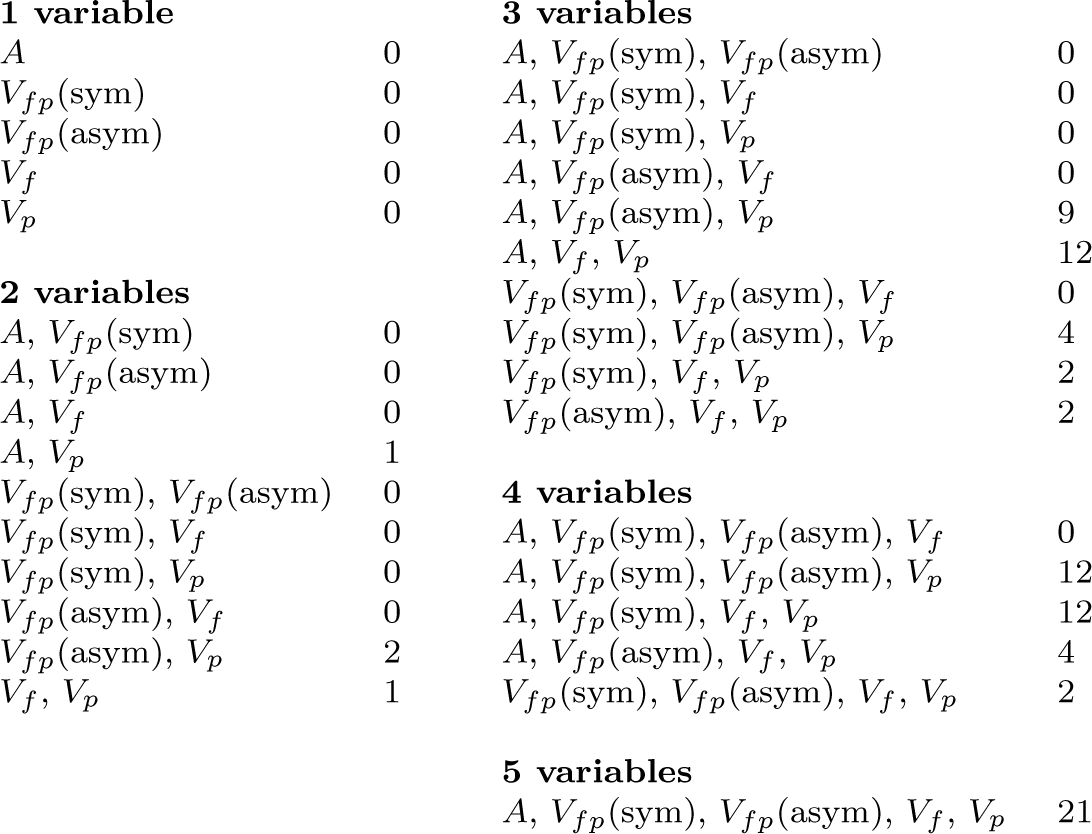
Number of preSMA neurons encoding sets of key task variables in best-fit GLM.

To capture the dynamics of the accumulating decision signal, we analyzed activity from 750 ms prior to 100 ms past arm movement initiation (Fig. 4A-C). We used demixed Principle Component Analysis (dPCA; [23]), a modification of Principal Component Analysis (PCA). Like standard PCA, it decomposes multi-dimensional data into components which capture an optimized amount of the variance in the data, but dPCA adds constraints to the optimization that also capture the dependence of the data on selected task variables, or factors. This allowed us to identify the average decision-related activity component in the preSMA population and to compare it with the predictions of one of the main theoretical constructs of decision making, the competing accumulator model [24].

**Fig. 4:**
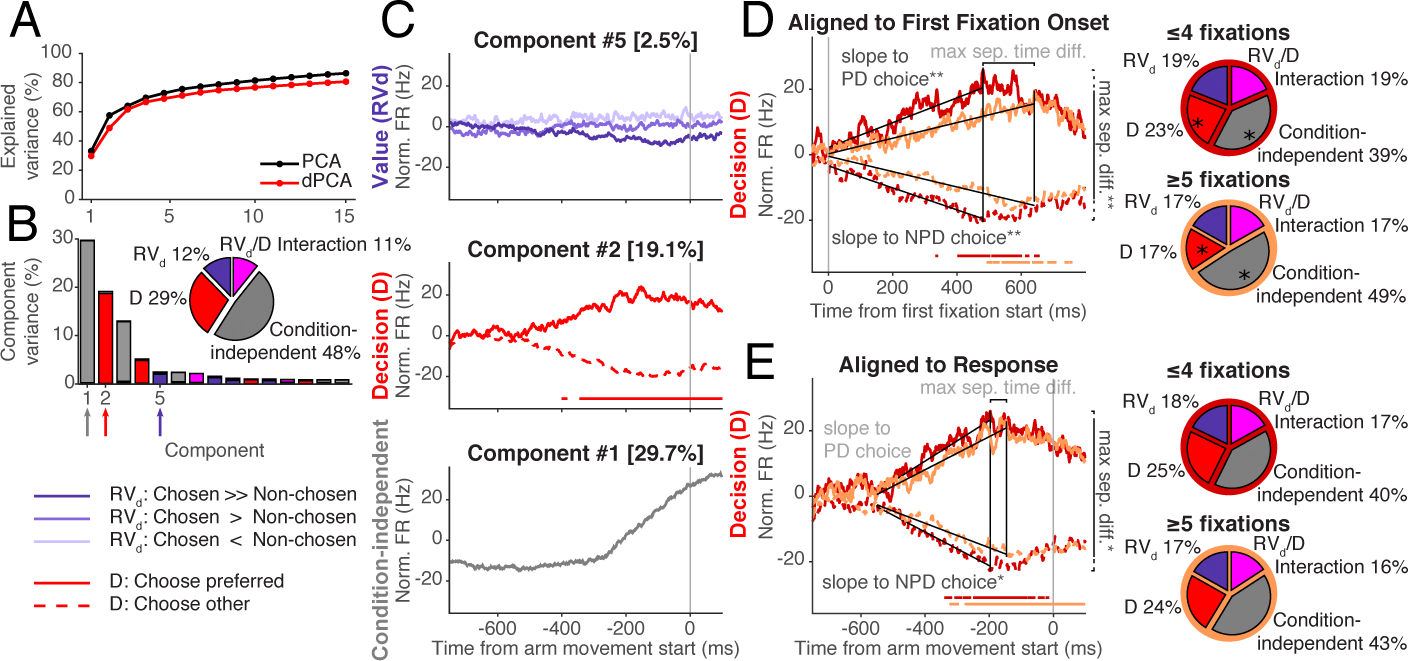
De-mixed PCA (dPCA) for neuronal activities preceding the choice. Neural activity aligned to the choice (i.e., the arm movement onset) was subjected to dPCA to identify the components related to two key factors: value (*RV_d_*, the difference between the subjective values of the chosen and non-chosen options) and decision (the position, preferred or non-preferred, of the chosen option), as well as the interaction of these factors and conditionindependent components. (A) Explained variance across components for dPCA and PCA. (B) Histogram of the first 15 components with regard to the value and decision factors, their interaction, and condition-independent variance. The pie chart shows the overall distribution of explained variance across these factors. Components 1, 2, and 5 (arrows under abscissa; colors correspond to those in pie chart and in panel C) are the largest condition-independent, decision-related, and value-related components, respectively. (C) Time course of the top-ranked value-related, decision-related, and condition-independent components, i.e. the projections of fixation-aligned neuronal spike densities onto the individual components. For Components 2 and 5, periods of significant classification (see the Classification Based on Demixed PCs section in Methods) are denoted with a horizontal line at the bottom of each plot. (D–E) Top-ranked decision components for dPCA on trials with fewer (*≤* 4; dark red lines) and more (*≤* 5; light red lines) fixations. Significant differences in total explained variance between these conditions for specific factors are denoted by *∗* on the pie charts. In D and E, periods of significant classification of the decision factor based on the plotted component are shown as horizontal lines in the same color as the corresponding time course plot. Also, component plots in D and E are annotated with four metrics that were computed and tested for significant differences between short and long trials: 1) the maximum separation between trials on which the PD option was chosen and trials on which the NPD option was chosen, 2) the time at which this maximum separation occurred, 3) the slope leading up to the point of maximum separation for trials when the PD option was chosen, and 4) the slope leading up to the point of maximum separation for trials when the NPD option was chosen. Significant differences in these metrics are denoted with *∗* (*p <* 0.05) or *∗∗* (*p <* 0.01). (D) dPCA for activity aligned to first fixation onset. (E) dPCA for activity aligned to arm movement onset.

We found that dPCA explained the variance of neuronal activities nearly as well as standard PCA (Fig. 4A). The first fifteen components are well de-mixed into three main factors (Fig. 4B). The components that explained most of the variance in neuronal activity for each of the factors are shown in Fig. 4C. The decision component (#2; orange) showed diverging activity patterns when the monkey chose the option in the PD *vs.* NPD location. The competing accumulator model predicts that decision-related activity gradually emerges over an extended time period. Indeed, the activity started to separate more than 500 ms before arm-movement onset and increased gradually until it reached a maximum about 200 ms before arm movement initiation.

The speed with which the decision-related component rises should be related to the length of the decision process [25–27]. To test this, we divided the trials into short inspection sequences (*≤* 4 fixations before choice) and long inspection sequences (*≥* 5 fixations before choice), and repeated the dPCA analysis separately for each. We first analysed neuronal activity starting from the beginning of the decision process, i.e. the first fixation, until 800 ms into the process of attribute inspection (Fig. 4D). The decision component rises faster during short than long inspections. The slope of the activity increase (when PD is chosen) and decrease (when NPD is chosen) are significantly different (both slopes: *p ≤* 0.004; permutation test, see the Permutation Tests of Demixed PCs section in Methods). Consequently, this component starts to dominate preSMA population activity earlier and explains a larger part of the total variance in short than long inspection sequences (Fig. 4D, pie charts: *≤* 4 fixations: 23%; *≥* 5 fixations: 17%; *p <* 0.001; permutation test). Thus, the speed with which the decision signal in preSMA grows predicts the length of the decision process.

The evaluation process should end when the difference in evidence has reached a critical threshold, at which time the monkey commits to one of the two options. Indeed, when aligned on movement onset (Fig. 4E), there was no significant difference in the time at which the decision component reached maximum difference for both short and long inspection sequences (*p >* 0.27; permutation test). There was also no difference in increasing slope to PD choices (*p >* 0.13; permutation test). The decreasing slopes to NPD choices were slightly, but significantly different (*p <* 0.049; permutation test). However, there was no significant overall difference in the strength of the decision component for short and long inspection sequences in the time period immediately preceding movement onset (23% and 24% for *≤* 4 and *≥* 5 fixations respectively; see Fig. 4E pie charts, *p >* 0.83; permutation test). Thus, the decision component of the preSMA population activity mostly fits the predictions of the competing accumulator model.

### 2.3 How Attention influences Action Value Signals

Our task requires the monkeys to sequentially fixate a series of spatial locations that provide decision-relevant information. These overt shifts of spatial attention provide a behavioral marker of which option and attribute is attended at each time during the decision-process. This allowed us to investigate the effects of attention on the decision process.

Three possible models of the mechanism by which attention could influence this process have been suggested, the additive, gain-modulation, and serial models. Each of these models predicts specific effects on neuronal activity (Fig. 1B,C,E). We identified preSMA neurons that matched these predicted effects, using a series of GLMs of neuronal activity during attribute inspections, within 50 ms after the beginning of each fixation until its end. These GLMs explained neuronal activity as a function of spatial attention (*A*) and a range of attention-dependent (*V_f_*, *V_fp_*) and -independent (*V_p_*) value signals (Fig. 5A).

**Fig. 5:**
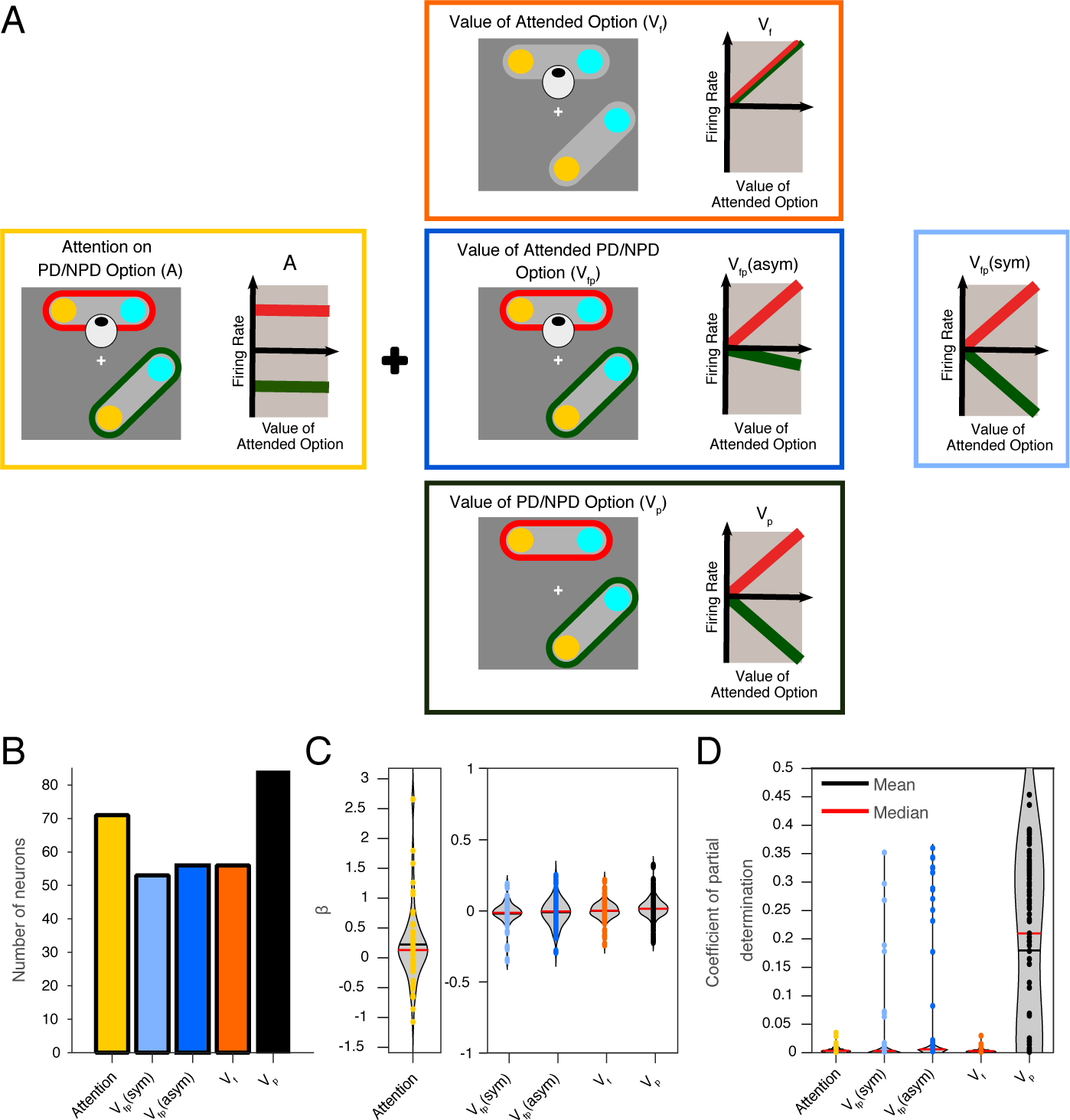
Generalized Linear Models (GLMs) for neuronal activities during fixation of attribute cues. (A) We analyzed the activity of preSMA neurons in the Attribute Search Period (50 ms after the beginning of each fixation to an attribute cue to its end) using GLM models consisting of a range of different functional signals predicted by the three models of attentional modulation. Each box contains a schematic depicting the dependency of each signal on the position of each option (indicated by the color border; red: PD, green: NPD) and whether or not it is attended (indicated by the eye). To the right is a schematic of the expected firing rate as a function of the value of the attended option (red: PD, green: NPD). Attention (A) indicates the spatial orientation of attention. The signal is binary, since attention is in either the PD or NPD. Value signals differ with respect to the framework for identifying the object they refer to. Such signals are predicted by the additive attention model (yellow box border). [Fig. 5 caption continues on next page.] Fixation-dependent value signals (*V_f_*) indicate the value of options based on whether they are attended or not (eye), independent of their position (no color borders for option symbols). Activity is expected to reflect value independent of whether the PD or NPD option is attended. Such signals are predicted by the serial attention model (orange box border). Fixation and position-dependent value signals (*V_fp_*; blue box borders) indicate the value of options based on whether a specific position (PD/NPD; color borders) is attended or not (eye). *V_fp_* signals come in two types. In an asymmetric *V_fp_* signal, the gain with which activity reflects value is different when the option in the PD or NPD position is attended and indicates that attention modulates the value signals. Such signals are predicted by the gain-modulation model of attention (dark blue box border). For symmetric *V_fp_*signals, the gain for the value representation is the same for PD and NPD options. This indicates simply the process of value accumulation, but no specific modulation by attention (light blue box border). Position-dependent value signals (*V_p_*; black) represent the value of options in specific positions (PD/NPD; color borders), independent of where attention is allocated (no eye). (B) The number of neurons whose best-fitting GLM includes specific predictor variable(s): Attention, *V_fp_*(symmetric), *V_fp_*(asymmetric), Vf, *V_p_*. The best model for each neuron was determined by comparing the AICs of models. The colors refer to the scheme used in A. (C) Coefficients *β* for each variable in the best-fit GLM for each neuron. (D) Coefficients of partial determination for each variable for all neurons.

The attention signal (*A*) indicates the spatial location of the currently fixated option (Fig. 5A; yellow box). This results in a binary shift of activity levels, as predicted by the ‘additive’ model of attentional modulation. A large number of preSMA neurons (A: 71/84, 85%) encoded the currently attended option, i.e., whether the fixated option is at the PD of the neuron (Fig. 5B). However, the attention signal was only encoded in conjunction with value-related signals (Table 1). This indicates that attention-location is not itself explicitly encoded, but rather modulates decision-related preSMA activity.

In our task, two basic reference frames could be used to determine the identity of the option whose value is encoded: (1) whether an option is fixated (*f*) or not, or (2) the spatial position (*p*) of an option in the visual display. This gives rise to three types of value signals (*V_f_, V_p_, V_fp_*) that reference option identity in three different ways (Fig. 5A).

First, we tested ‘position-dependent’ value signals (*V_p_*) that represent the value of the option located in the neuron’s PD, irrespective of where attention was directed (Fig. 5A, black box). These signals represent the total currently accumulated evidence about a specific option continuously throughout the decision process. Signals of this type are predicted by standard accumulator decision models. In contrast, the ‘serial’ model strongly predicts that position-dependent value signals should not exist. Our findings were unequivocal. All 84 preSMA neurons encoded position-dependent value signals (Fig. 5B and Table 1). Thus, preSMA neurons robustly represented the value of their PD option throughout the entire sequence of fixations. This strongly indicates that attentional modulation of action value signals in the preSMA is not so strong that it leads to a serial decision making process. Instead, the decision-process seems to operate by evaluating both options in parallel. However, nearly all neurons (83/84, 99%) also reflected value signals that depended on which option was currently attended. Thus, the decision-related activity in preSMA is modified by sequential shifts in attention.

Second, we tested ‘fixation-dependent’ value signals (*V_f_*). These represent the value of the currently attended option, irrespective of its spatial location (Fig. 5A, orange box). We found a large number (56/84; 67%) of preSMA neurons carrying such signals (Fig. 5B). Because serial models of attentional modulation assume that only the attended option is represented, they predict such signals. However, all fixation-dependent signals (*V_f_*) were carried by neurons that also carried position-dependent signals (*V_p_*; Table 1). The values of the coefficients *β* (in the best-fit model) and the coefficient of partial determination (see the Generalized Linear Model section in Methods; Eq. 8) of *V_f_* signals were not stronger than those of other value signals (Fig. 5C,D). These facts argue against a ‘serial’ model of attentional modulation. An alternative interpretation is that the *V_f_* signal represents a more general decision- or motor-related process, such as arousal.

The third type of value signals we tested were ‘fixationand positiondependent’ value signals (*V_fp_*) that combine both the fixationand positiondependent reference frame. They represent the value of the currently attended option contingent on whether it is in the neuron’s PD or not (Fig. 5A, blue box).

We distinguished two patterns of activity. One pattern was symmetric across the positions, so that any value-related activity for the PD option was matched by an equally strong, but inverse, activity for the NPD option (Fig. 5A, light blue box). This signal reflects information about the value of both options, but in a manner that reflects a comparative evaluation process, whereby the preSMA neurons encode the expected value of an arm movement in their PD, relative to the expected value of the NPD movement. Thus, this activity pattern is a product of the decision process as driven by sequentially attended information, not an indication of attentional modulation. In line with the basic role of this signal in the decision process, many preSMA neurons (53/84; 63%) carried symmetric *V_fp_* signals (Fig. 5B).

In contrast, the other pattern was asymmetric across the positions (Fig. 5A, dark blue box). These signals reflected value, but with different strengths for the PD and NPD positions. Thus, these signals reflected the value signal stronger when specific options were attended. Such signals are predicted by the ‘gain-modulation’ model of attentional modulation. A large number of preSMA neurons (56/84; 67%) encoded asymmetric value signals. This strongly supports the ‘gain-modulation’ model.

In conclusion, all tested variables were strongly represented in the preSMA (Fig. 5B). All neurons encoded more than one variable and the majority of neurons (60%; 51/84) encoded all or all-but-one of the variables (Table 1). Neurons encoding all or most variables were selected as the best fit models for neurons far more often than models encoding fewer variables (*p <* 1.7 *×* 10*^−^*^11^; *χ*^2^ goodness-of-fit test). Thus, the variables are best understood not as functionally independent signals, but rather as a decomposition of a complex signal carried by many neurons. This complex signal seems to be an attention-independent value signal that indicates the value of an option in a particular position continuously throughout the inspection sequence, which is modulated by two time-varying, attention-dependent signals, one additive, the other multiplicative.

### 2.4 Temporal Dynamics of Attentional Modulation

To better understand the time dependence of attentional modulation of the decision process, we applied dPCA to preSMA neuronal activity during the attribute fixation period (Fig. 6). The dPCA explains the variance of neuronal activities very well (Fig. 6A). The activity pattern captured by the first component explains a very large part of the total variance (34.1%), and shows a mixture of three main effects (Fig. 6C).

**Fig. 6:**
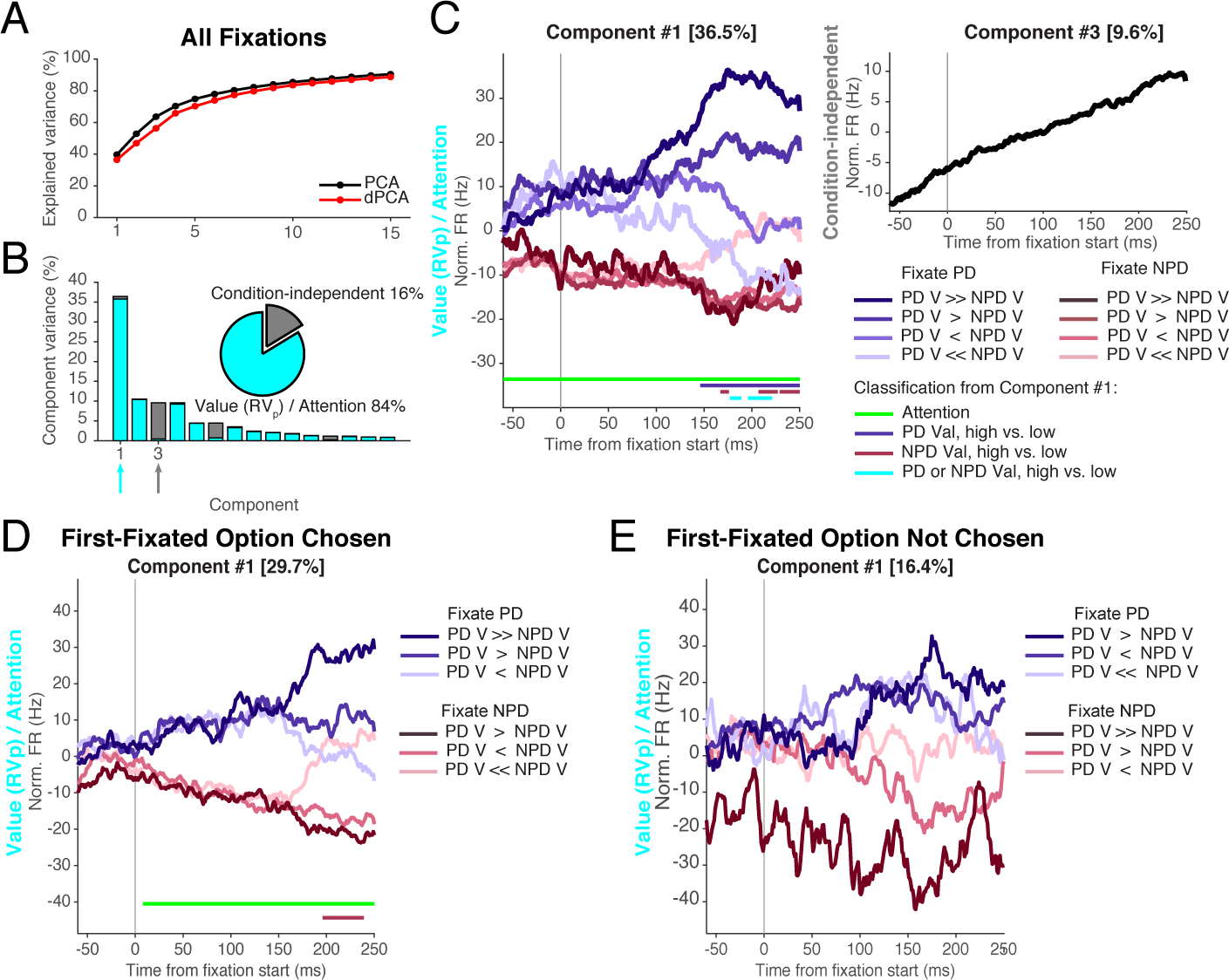
dPCA of Neural Activity during all fixations in the attribute search period. We demix activity using only one factor, called *RV_p_/A* (plus the condition-independent factor) which consists of the relative value *RV_p_* multiplexed with position-defined attention *A*. This factor has eight states, see legend under panel B. (A) Explained variance across components for dPCA and PCA. (B) Histogram of the first 15 components with regard to the *RV_p_/A* factor and condition-independent variance. The pie chart shows the overall distribution of explained variance across these factors. Components 1 and 3 (arrows under abscissa; colors correspond to those in pie chart) are the largest components of the *RV_p_/A* factor and the condition-independent activity, respectively. (C) Upper panel: Time course of the eight states of *RV_p_/A*, i.e. the projections of fixation-aligned neuronal spike densities onto the individual components. For Component 1, periods of significant classification (see the Classification Based on Demixed PCs section in Methods) are denoted with a horizontal line at the bottom of each plot, see legend for color coding. Where no horizontal line is shown, the state could not be significantly classified based on component 1. Lower panel: Time course of projection onto the condition-independent factor. (D,E) Same as in C but only for first fixations in each trial: (D) Trials in which the first fixated option was chosen, (E) Trials in which the first fixated option was not chosen. Here reduced sets of states for the *RV_p_/A* factor are used, see legends under panels D and E.

First, activity is higher when the PD option is fixated (blue lines) than when the NPD option is fixated (red lines). This reflects the ‘additive’ effect of attention captured by the *A* variable in the GLM analysis (Fig. 5A, indicated in yellow). The effect of attention on neuronal activity varied only moderately during the fixation period. Using component #1 allows us to classify which option was attended for nearly the entire time analyzed (Fig. 5C, green line). When attention switched to a new option (first fixation), the additive modulation started at the very beginning of the fixation, but was not present before fixation onset (Extended Data Fig. 3 A,C). In contrast, when attention stayed within the same option (Extended Fig. 3 B,D), the additive effect precedes the fixation and is present throughout. This is a strong indication that it reflects attentional state and not a response to a new visual stimulus or eye position. Second, value influenced activity starting about 100 ms after fixation onset (divergent blue lines). The delay likely reflects the latency until fixated visual information could influence the decision process in preSMA, due to preceding visual and value estimation processing. Importantly, the value-related activity modulation overall was much stronger when the PD option was attended (blue lines) than when the NPD option was attended (red lines). This type of attention-modulated value response reflects a gain-modulating effect of attention on preSMA activity captured by the asymmetric *RV_fp_* variable in the GLM analysis (Fig. 5A, indicated in dark blue). We can classify value significantly above chance later in the analyses window for high *vs.* low value when considering either PD *or* NPD options (violet and red lines).

Third, high value leads to increased activity, even if the NPD option is attended (see lightest red line). This effect coexists with the gain-modulation effect that in all other cases suppresses value-related activation for NPD options. This reflection of the attended option value regardless of its position is captured by the fixation-dependent value signals (*V_f_*) in the GLM analysis (Fig. 5A, indicated in orange).

For value, we consider three binary classifications: high *vs.* low when the PD option is fixated, high *vs.* low when the NPD option is fixated, and high *vs.* low regardless of what is fixated. Finally, the condition-independent component (#3) captures a general increase in firing that occurs throughout the decision process, here observed over the duration of single fixations. This slowly rising activity component in the preSMA activity might reflect an urgency signal ([28–30]; but see [31]).

To test if the strength of the attentional modulation of the action value signals had an effect on decision-making, we repeated the dPCA for fixations to the first inspected option separately for trials in which the monkeys eventually chose this option *vs.* those where they ended up choosing the second inspected option (Fig. 6D,E). The largest component shows clear differences. Both the additive and gain-modulation effects were substantially stronger when the monkeys chose the attended option (Fig. 6D). In contrast, the effects were noticeably weaker when the attended option was not chosen (Fig. 6E). Repeating the classification procedure, we see classification significantly above chance for attention and value of the NPD option only for first-fixated options that were chosen. This suggests that the attention-related modulation increases the probability that the attended option will be chosen.

## 3 Discussion

Decisions among multiple options that differ across multiple attributes are common, but complex, decision problems. Naturalistic choices typically involve the sequential engagement of attention to different options and their attributes to gather information before a choice is made. Primate electrophysiology experiments have started to investigate the neuronal circuits involved in these attention-guided decisions [6, 8, 9]. However, many aspects of the underlying neuronal mechanisms by which attention influences decision-making are still not well understood. In most classical value-based decision tasks, shifts of attention are assumed to be covert and essentially unknowable. In contrast, in our task the free viewing condition allows us to observe the shifts of spatial attention between different attributes and options, and to directly observe the effect of attention on decision-related neuronal representations.

PreSMA activity reflects the dynamics of the decision process and allows insights into the functional architecture of the network underlying the decision process. This holds independent of whether the preSMA plays a causal role in the decision process, or just reflects the momentary state of the ongoing decision process. This allows us to investigate three mechanistic hypotheses about the role of attention in the decision process: the additive, gain-modulation, and serial models of attentional modulation (Fig. 1). Our findings support the additive and gain-modulation models, but argue against the serial model.

Our results indicate a basic parallel decision architecture (Fig. 1A). The value of each action is represented by an independent pool of neurons (each labeled by its preferred spatial direction). Pools compete through mutual inhibition [24, 32]. Activation strength of each pool is primarily a function of the information about the value of its respective option. This value input likely comes from areas outside of the premotor network, such as orbitofrontal or ventromedial prefrontal cortex.

Another network controls the current location of attention. Input from this network has two effects. First, it directly enhances activity of the preSMA pool representing the currently attended option and suppresses activity in pools representing unattended options (Fig. 1C). This firing rate modulation is constant and independent of the value difference of the two options (Fig. 5A, yellow). This finding supports the ‘additive value’ model suggesting that attention to a response option directly increases its value and thereby increases the preference for the attended option [33–35]. In line with this prediction, the strength of the additive modulation in preSMA is correlated with the likelihood that the attended option is chosen (Fig. 6F,I). This effect could explain behavioral findings showing that manipulating attention changes choice [5, 19, 20, 36–38], and our finding that the length with which the monkeys attended an option is correlated with the probability that they choose it (Fig. 2E).

Second, attention also leads to a gain-modulation of the preSMA responses to the currently attended attribute information (Fig. 5A, blue). It is known that attention influences the strength and acuity of sensory representations [39–43]. Similar attention-driven activity modulations have been reported for option value signals in the orbitofrontal cortex [6–9]. Attended information should therefore have a stronger influence on decision-making than unattended information. In contrast, the preSMA action value signals change very little, when the non-preferred option is attended (Fig. 6C). Thus, attention seems to partially shield the unattended option from the competition of the attended one. This could counterbalance the ‘additive’ attention effect and might lead to a more balanced evaluation of both options, particularly in tasks like ours that requires serial shifts of attention to sample the decision-relevant information. It has been suggested that attentional effects are so strong that only the population that encodes the value of the currently attended option is active, while the other one is suppressed [13, 14]. This would result in a serial evaluation of the reward options in a sequence of accept/reject choices of the currently attended option (i.e., the ‘serial’ model). We have found evidence for fixation-dependent value signals (Fig. 5A, orange). Nevertheless, the preSMA neurons also carry value signals that indicate the accumulated value estimates for both options, independent of which option is attended (Fig. 5B). This finding argues very strongly against the ‘serial attention’ model.

While our results rule out a strictly serial decision architecture, they are compatible with a partially sequential architecture, in which only the currently attended and the currently best option are represented [6]. This partially sequential architecture would suggest a functional role for the additive attentional effect, because it would ensure that attended options are able to successfully compete in the face of strong inhibitory competition which otherwise would not allow the attended option to overcome the inhibition. In a completely parallel architecture the enhancement effect would only lead to distortions of the decision process. Distinguishing between the partially sequential and the completely parallel architectures requires experiments with more than two options.

In conclusion, our findings show that attention influences decisions through an additive and a gain-modulation mechanism. Together, these modulations select the type of information on which the decision is based and in this way can significantly influence the final choice.

## Supplementary information

See supplementary information for further details on modeling of the monkeys’ choice behavior, discussion of the evidence for their understanding of the symbolic cues, and an illustration of the definition and binning of value, attention, and decision factors for dPCA.

## Declarations

- Funding: Supported through NSF grant 1835202, NIH grant R01DA040990 (CRCNS), Office of Naval Research N00014-22-1-2699 and NIH Fellowship 5K00NS105204-05
- Competing interests: The authors declare no competing interests.
- Ethics approval: All animal care and experimental procedures were conducted in accordance with the US public Health Service policy on the humane care and use of laboratory animals and were approved by the Johns Hopkins University Institutional Animal Care and Use Committee (IACUC).
- Data and code availability: Data and code are available through the Johns Hopkins Research Data Repository.
- Authors’ contributions: M.U., D.L., E.N., and V.S. designed the behavioral task. Y-P.Y. collected the data. A.S. and Y-P.Y. analyzed the data. A.S., Y-P.Y., E.N. and V.S. wrote the manuscript.

## 4 Methods

### 4.1 Experimental Procedures

Two male rhesus monkeys (Monkey I, 8 kg, 10 years; Monkey H, 11 kg, 10 years) were used in this study. Each animal was implanted with a head-restraint device including a recording chamber over the medial frontal cortex with access to the supplementary motor area (SMA) and pre-SMA. All animal care and experimental procedures were conducted in accordance with the US public Health Service policy on the humane care and use of laboratory animals and were approved by the Johns Hopkins University Institutional Animal Care and Use Committee (IACUC). During the experimental sessions, the monkey was seated in an electrically insulated enclosure with its head restrained, facing a video monitor. MonkeyLogic software (https://www.brown.edu/Research/monkeylogic/) was used to control task events, stimuli, and reward, as well as monitor and store behavioral events [44–46]. Eye positions were monitored with an infrared corneal reflection system, EyeLink 1000 (SR Research, Kanata, ON, Canada) at a sampling rate of 1000 Hz. All analyses were performed using custom Matlab (Mathworks, Natick MA) code, unless noted otherwise.

### 4.2 Behavioral Task

Two monkeys were trained to perform a sequential sampling decision-making task (Fig. 2A) in which they were required to choose between two risky options. A trial started with the presentation of a fixation cue (‘+’) in the center of the screen. Monkeys were required to fixate this cue for 100-400 ms. Following fixation, two options, each with two attributes (reward amount and reward probability) were presented on the screen with a specific spatial configuration drawn randomly from three possible configurations (Fig. 2B). Attributes belonging to the same option were connected by a light gray bar. Magnitudes of the two attributes, reward amount and reward probability, were represented by horizontal and vertical tags and covered by yellow and blue masks, respectively (Fig. 2C). There were 4 levels of reward magnitude for monkey I ([0,1,2,4] water units) and 5 levels for monkey H ([0,1,2,3,4] water units). Likewise, there were 4 levels of reward probability (P(win) = [0,0.25,0.5,1] for monkey I and 5 levels for monkey H (P(win) = [0,0.25,0.5,0.75,1]). Locations of the two attributes within a option were randomized. The magnitude of each attribute was revealed only when the animal fixated on its location, which removed the colored mask and unmasked the attribute. Fixation detection and unmasking only took *≈* 20ms, resulting in a nearly natural viewing experience.

Subjects were free to visually inspect the information of each attribute as much/long as they wanted to and ultimately indicated their choice by moving a joystick to the location that corresponded to the chosen option. The monkeys would initially hold the joystick in the middle position and could move it in these three directions only: left, right and up. The timing, direction, and amplitudes of joystick movement and eye movements were recorded. Following choice, the option that was *not* chosen disappeared and only the chosen option stayed on the screen for 200 ms (Feedback Epoch). Then the gray bar connecting the two attributes within the chosen option became darker during the immediately following 200 ms (Reward Epoch) and a water reward was delivered or not, depending on the drawing of the random variable controlled by P(win). Trials were separated by a 2000 ms inter-trial interval. Aborted and error trials were followed by a 1500ms punishment time period and were presented again later.

Trials were selected from 1 of 2 types: non-zero-option trials (80%) and zero-option trials (20%). The non-zero-option trial consisted of two non-zero options (options with no attribute equal to zero)while the zero-option trial consisted of one zero option (option with an attribute equal to zero)and one non-zero option. Trials were presented in a block design: Blocks containing all 120 possible non-zero-options in pseudo-random order alternated with blocks of 30 zero-option trials, pseudo-randomly drawn from all possible zero-option trials.

The two monkeys experienced extensive (*>* 30) training sessions before collecting their behavioral data with neuronal recording. We included behavioral data from Monkey I performing 10 sessions (585 *±* 84 completed choice trials) and Monkey H performing 20 sessions (799 *±* 61 completed choice trials) with corresponding neural data in this study.

To assess how the sequence of fixations related to choice behavior, fixates were separated according to the order in which they occurred (i.e. first, second, etc.) and the proportion of trials on which the currently fixated option would be chosen was computed for each group. Significance of these choice probabilities (Fig. 2I) was assessed for each fixation in sequence using a t-test of the null hypothesis that the proportion of trials on which the currently fixated option would be chosen comes from a distribution with mean 0.5.

### 4.3 Behavioral Modeling of Monkeys’ Risky Choices

There are several ways in which the monkeys could have integrated reward amount and probability information to estimate the overall value of an option. We considered four possible models. For all, only the (unordered) *set* of fixation targets was taken into account, not their (ordered) *sequence*.

The first model is the expected value (EV) model which assumes that monkeys calculated the value of an option by finding the product of amount (*a*) and probability (*p*):

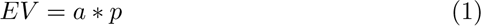

where *a* and *p* are the reward amount and reward probability of an option, respectively. The EV model thus is based on the assumption that monkeys can compute the linear combination of *a* and *p* with a multiplication rule.

Second is the prospect theory (PT) model which assumes monkeys again integrate *a* and *p* with a multiplicative rule but with specific nonlinear functions that transform attributes *a* and *p* to their corresponding subjective mappings, i.e. utility *u*(*a*) and weighted probability *w*(*p*) [47]. Utility is defined as:

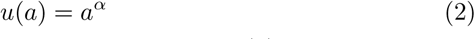

where *α* is a parameter in the nonlinear mapping of *a* to *u*(*a*). A utility function with 0 *< α <* 1 results in a concave curve, indicating the monkey underweights high amounts while *α >* 1 gives a convex utility function, indicating that the monkey over weights high amounts.

The weighted probability is computed as,

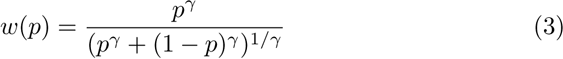

where *γ* is a parameter in the nonlinear mapping of probability *p* to weighted probability *w*(*p*). A probability function with 0 *< γ <* 1 results in an Sshape curve, indicating the monkey underestimates lower probabilities while overestimating higher probabilities. A probability function with *γ >* 1 gives an inverse S-shape curve, indicating the monkey overestimates lower probabilities while underestimating higher probabilities.

The PT model calculates the value of an option as the product of *u*(*a*) and

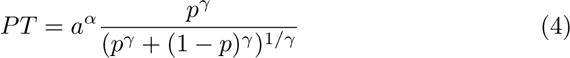

The crucial difference between EV and PT is that the PT model assumes a nonlinear (subjective) mapping of *a* and *p*. For the special case *α* = *γ* = 1, PT becomes EV.

Third, the additive value (AV) model assumes that monkeys calculate the value of an option by adding the rank of its corresponding amount and probability attributes (range 1–5 for both):

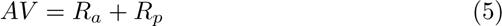

where *R_a_* and *R_p_* are the ranks of the amount and probability attributes respectively. Note that for Monkey I amount values of 3 and probability values of 0.75 were not used, and therefore *R_a_* = 4 and *R_p_* = 4 do not occur.

Fourth and final is the weighted additive value (WAV) model, a generalization of the AV model. As in the AV model, we assume that monkeys integrate the ranks of attributes to obtain an option value using an additive rule. However, in the WAV model one of the two attributes may have a different importance/significance than the other. The relative weight is implemented by introducing a free parameter *w_a_*:

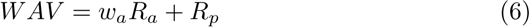

where 0 *< w_a_ <* 1 means the monkey weighs amount less than probability, indicating risk-averse behavior. On the other hand, *w_a_ >* 1 means the monkey weighs amount more than probability, indicating risk-seeking behavior.

For models with at least one free parameter (PT and WAV), parameters were set based on the Bayesian Information Criterion (BIC, a measure of model fit taking into account both agreement of empirical data with model predictions and the number of free parameters [48]):

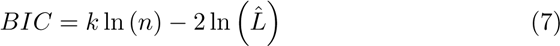

where *k*is the number of free parameters in the model, *n* is the number of observations used in fitting the model, and *L*^^^ is the likelihood of the model predicting the animal’s choice. Higher prediction accuracy indicates a better fit, and lower BIC values indicates that a better compromise of a good fit is obtained with a small number of free parameters.

All trials from all recording sessions when the monkey fixated all four unique attributes and then made a choice were used to set the parameters, separately for each monkey. In Supplementary Table S1, we compare the prediction accuracy (percent agreement between the animals’ choices and the choices that are predicted by the models) and BIC for all four models and both monkeys. Based on BIC, the WAV model was used for all analyses of value-related neural activity, but analyses were repeated using PT (see Extended Data).

### 4.4 Saccade Detection and Definition of Choice Events

Eye movements were classified offline (after recordings were completed) using a computer algorithm that searched first for significantly elevated eye velocity (*>* 30*^◦^*/s). Saccade initiation was defined as the beginning of a monotonic change in eye position starting 15 ms before the high-velocity gaze shift. A choice event was defined to have occurred if a joystick movement started in the center (1*^◦^* x 1*^◦^*) and ended in one of the peripheral target windows (2.5*^◦^* x 2.5*^◦^*).

### 4.5 Fixation Timing

During the experiment, fixation onset was detected on-line and used to control the presentation of the visual stimuli. However, when a fixation was detected, the monkey already had started to fixate a particular attribute. In order to define the actual behavioral fixation time windows, we analyzed the eye traces off-line. We compared tracked eye position data with millisecond resolution to the positions of options presented on the screen and the time of their unmasking (triggered by real-time fixation detection). This allowed us to determine the precise timing of the fixations the animal made to each attribute. For each recorded unmasking of an attribute, the fixation is defined as the period extending forward and backward in time, during which the eye position was stable and directed toward the unmasked attributed. Eye position was considered stable when the six-point moving average of the Euclidean distance between successive time points of the tracked eye position on the screen was less than 2.3 standard deviations above the average distance between position at successive time points over the course of the trial. This empirically set threshold captures the vast majority of saccades between fixations. For the rare cases where it does not (i.e. in which multiple unmasking events occur between threshold crossings), the fixations are delineated by the maximum of the smoothed point-to-point distance between the unmasking events.

### 4.6 Neurophysiological Recording

Single tungsten microelectrodes (impedance of 2-4 MΩ, Frederick Haer, Bowdoinham, ME) were used for extracellular single neuron recording. Electrodes were inserted through a guide tube positioned just above the surface of the dura mater and were lowered into the cortex under control of a custom-built microdrive system. The electrodes penetrated the brain perpendicular to the surface of the cortex. The depths of the neurons were estimated by depth relative to the surface of the cortex. Recorded signals from the electrodes were amplified and bandpass filtered before action potentials were detected by a time-amplitude window discriminator. Spikes were isolated online if the amplitude of the action potential was sufficiently above a background threshold to reliably trigger a time-amplitude window discriminator and the waveform of the action potential was invariant and sustained throughout the experimental recording. Spikes were then identified using principal component analysis (PCA) and the time stamps were collected at a sampling rate of 1,000 Hz (Plexon, Dallas TX).

We recorded neurons from the medial frontal cortex (including SMA and pre-SMA). Recording sites were reconstructed with T1 and T2 magnetic resonance images (MRIs) obtained for each monkey at 3 Tesla. We used the known stereoscopic recording chamber location and recording depths of the electrodes to estimate the location of each recorded neuron. The estimated recording locations were superimposed on the MRI scans of each monkey, see Supplementary Fig. S1. Cortical areas were estimated of the macaque monkey brain atlas by [49].

### 4.7 Preferred Direction of Neurons

For every neuron a preferred direction (PD) was defined as the direction of arm movement indicating choice (to top, right, or left) that corresponded to the highest mean firing rate across all trials for that neuron. For most, but not all neurons, there was a statistically significant difference in firing rate based on a one-way ANOVA to test the null hypothesis that movements to all three directions had the same mean firing rate.

### 4.8 Generalized Linear Model

To quantitatively characterize the variables that each neuron encodes during the Response Period, defined as the period from 200 ms before to joystick movement onset, we fit a series of generalized linear models (GLMs). We tested if the neuronal activity was related to the following variables: decision (*D*), value of the chosen option (*V_d_*), position-dependent value of the chosen option (*V_dp_*), relative value of the chosen option (*RV_d_*), and position-dependent relative value of the chosen option (*RV_dp_*). The Decision variable (*D*) is binary (+1 for PD and -1 for NPD) and represents whether the selected arm movement is directed towards the neuron’s preferred direction (PD) or the non-preferred direction (NPD). The value variable (*V_d_*) encodes the value of the chosen option independent of whether the PD or NPD option is chosen. In contrast, the position-dependent variables do depend on whether the PD or NPD option is chosen; there are three forms these variables (*V_dp_*) . The first form, *V_dp,P_* _+*NP*_, encodes the value of the option with positive sign when the option is in the PD and with negative sign when it is in the NPD. The second form, *V_dp,P_*, encodes the value of the chosen option if it is in the PD; if it is in the NPD the variable is set to zero. The third form, *V_dp,NP_*, is the inverse: its value is set to the chosen value if the option is in the NPD and to zero if it is in the PD. Relative value variables follow the same structure. The relative value variable (*RV_d_*) encodes the value difference between the chosen and non-chosen option independent of their position. There are again three forms of position-dependent relative value (*RV_dp_*) variables. The first form, *RV_dp,P_* _+*NP*_, encodes the relative value between options with positive sign if the chosen option is in the PD and with negative sign if it is in the NPD. The second, *RV_dp,P_*, encodes the relative value of the chosen option when it is in the PD, and zero when it is in the NPD. The third, *RV_dp,NP_*, is its inverse: it encodes the relative value of the chosen option when it is in the NPD, and zero when it is in the PD.

We examined the activity of each neuron using a series of generalized linear models (GLMs) using logarithmic link functions, with the mean firing rate (FR) within the Response Period (200 ms preceding the arm movement) for each trial as the dependent variable, and predictors derived from the decision-related variables as the independent variables:

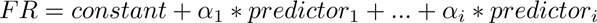

We tested all potential combinations of 5 types of basic variables (decision, decision-dependent value, decision- and preference-dependent value, decision-dependent relative value, and decision- and preference-dependent relative value) and a baseline term.

There are 9 basic variables: 1 encoding the decision, 1 the position-independent value of the chosen option, 3 encoding the position-dependent value of the chosen option, 1 encoding the position-independent relative value of the chosen option, and 3 encoding the position-dependent relative value of the chosen option. Using all possible combinations would allow us to construct a total of 2^9^ = 512 different models for each neuron. However, our data does not allow a robust differentiation of such a large number of models. We therefore constructed all models in which either zero or one of each *form* of variables was included. For instance, the position-dependent value of the chosen option form could only be represented by one variable (e.g. a model could include only *V_dp,P_* _+*NP*_, *V_dp,P_*, *OR V_dp,NP_*), but not combinations of these specific variables (e.g., *V_dp,P_* _+*NP*_ *AND V_dp,P_* or all three variables). Restricting the combinations of decision variables resulted in a total of 9 five-variable encoding models, 33 four-variable models, 46 three-variable models, 27 two-variables models, 9 single variable models, and one model containing only a baseline term. Thus, in total, we tested 125 models for each neuron. We determined the best-fitting model for each neuron using the Akaike information criterion (AIC) [50] as *AIC* = 2 *∗ df −* 2 *∗ LL_max_* where *df* is the number of variables in the model and *LL_max_* is the maximum log-likelihood of the GLM fit.

For the Attribute Search Period, we fit a series of GLMs as we did in the Response Period. Now, for each neuron the dependent variable was the mean firing rate during the period from 50 ms after the beginning of a fixation to the end of a fixation (i.e. the beginning of the following saccade), aligned on all fixations for each trial.

There are 11 fixation-dependent variables including their interactions: attention (*A*), attribute rank of amount or probability (*RankA* and *RankP*), value of fixated option (*V_f_*), position-dependent value of fixated option (*V_fp_* divided into *V_fp,P_* _+_*_NP_*, *V_fp,P_*, *V_fp,NP_*), relative value of fixated option (*RV_f_*), and position-dependent relative value of fixated option (*RV_fp_* divided into *RV_fp,P_* _+_*_NP_*, *RV_fp,P_*, *RV_fp,NP_*) and 2 fixation-independent variables: value of option in PD (*V_p_*) and relative value of option in PD (*RV_p_*).

The attention variable represents whether the fixated target was in the PD or NPD, indicated as 1 for PD and -1 for NPD. The fixated value (*V_f_*) and relative value (*RV_f_*) variables encode the value of the fixated option.

There are the same three interaction forms as during the Response Period but they are now defined with respect to the fixated option, rather than the chosen option. For instance, the variable *V_fp,P_* _+*NP*_ takes the value of the option with a positive sign when the PD is fixated, and with a negative sign when the NPD is fixated. As in the Response Period, not all possible 2^16^ = 65, 536 models were considered. Models with fixation- and preference-dependent value variables were considered in two categories, symmetric and asymmetric. Symmetric models contained *only V_fp,P_* _+*NP*_ while asymmetric models contained *both V_fp,P_* and *V_fp,NP_* allowing a separate coefficient for each (hence asymmetric representation). A given model could contain up to one each of the variables of the types *V_f_* and *V_fp_*, but each could be alternately defined as attribute rank (*RankA*/*RankP*), single option value, or relative value (difference in value between options). Similarly, a given model could contain only one variable of the type *V_p_*, but this variable could be defined alternately as attribute rank (*RankA*/*RankP*), single option value, or relative value (*RV_p_*). The rank of amount and probability took values 1–4 for monkey I and 1–5 for monkey H. As in the Response Period, we determined the best-fitting model for each neuron using the Akaike information criterion.

As a measure of the independent contributions of the variables in each best-fit GLM, we also compute the coefficient of partial determination (CPD) for each variable:

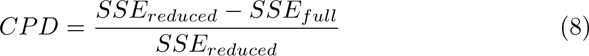

where *SSE_full_*is the sum of squared errors for the best-fit GLM and *SSE_reduced_*is the sum of squared errors for a model that contains all of the same terms *except* the variable for which the CPD is being computed.

### 4.9 Demixed Principal Components Analysis

A common representation of the population activity of *N* neurons is the Euclidean space spanned by *N* orthogonal axes, each of which corresponding to the activity of one of the neurons. Of interest is frequently the mean activity (averaged over trials) of the population as a function of time, e.g. the Peri-Stimulus Time Histogram or its equivalent for other time periods, like those aligned on motor responses. It is represented in this space as a time-dependent trajectory where each point corresponds to the set of average firing rates of one of the neurons. A commonly used method to reduce the dimension of this “point cloud” is Principal Component Analysis (PCA). Its components are a set of *N* orthogonal axes in this space defined as linear combinations of the original axes. The weights to form these linear combinations are chosen to maximize the amount of explained variance of the point cloud, with the amount of explained variance decreasing monotonically with increasing order of principal components.

While this is a useful technique in many cases, a common difficulty in the interpretation of neuronal activity, in particular in more central areas, is that multiple stimulus and internal state variables are represented in individual neurons. Such simultaneous coding for multiple variables by single neurons, called “mixed selectivity,” makes it difficult to understand the function of the whole circuitry and how it represents the task variables and internal states of the nervous system. Kobak et al. [23] introduced a generalization of PCA in which principal components are chosen not strictly to maximize explained variance but with the additional constraint that components, chosen at the outset of the analysis, attempt to separate (demix) stimulus and state parameters. In this “demixed PCA” (dPCA), the new axes are not necessarily orthogonal and therefore the explained variance is in general lower than in PCA. However, the projections of the population activity onto the new axes correspond to the contributions of the pre-defined experimental variables which makes their interpretations much more useful than that of PCA components where these contributions are mixed. We used the Matlab version of the dPCA code provided at http://github.com/machenslab/dPCA.

For all recorded neurons from both animals, spike density was first computed for each individual trial by convolving each spike with a one-sided (forward in time) exponential decay kernel (*τ* = 20 ms) to generate a time series sampled at 1 kHz. For the response-aligned analysis (Fig. 4), spike density time series (beginning 750 ms prior to the beginning of the choice-indicating arm movement and continuing 100 ms after the beginning of the arm movement) were placed in a four-dimensional array, with data organized according to neuron, the relative value of the chosen option (sorted into three bins), chosen direction (PD or NPD), and trial. For each neuron, spike densities were then averaged across all trials with identical decision and value. The resulting 87 neuron-by-3 relative value bin-by-2 directions (chosen)-by-850 ms array was subjected to dPCA, with the resulting de-mixed PCs labelled as being related to decision, value, the interaction of value and decision, or as being condition-independent.

Using this approach, there were sufficient data for all 84 neurons to be included in all analyses shown in Fig. 4. The total of all 61,862 trials was included for the analysis of all responses (Fig. 4A–C). For the analyses of trials with four or fewer fixations or five or more fixations, 27,645 and 34,217 trials were included respectively (Fig. 4D–E).

For the fixation-aligned analysis, spike density time series (beginning 50 ms prior to the beginning of each fixation and extending to 200 ms after the beginning of each fixation) were sorted into an array according to neuron, relative value of the fixated option (sorted into four bins), and position (preferred or non-preferred) of the fixated option. Spike densities were then averaged across all fixations (Fig. 6A–C) or for selected fixations (Fig. 6D–E and Extended Data Fig. 3) from all neurons with identical value and attention state. The resulting 87 neuron-by-3 or 4 relative value bin-by-2 direction (attended)-by250 ms array was subjected to dPCA, with the resulting de-mixed PCs labelled as being related to value, attention, an interaction of value and attention, or as being condition-independent.

Because this approach requires data in all attention and value states for all neurons, some neurons had to be excluded from analyses of specific subsets of fixations. For the analysis of all fixations (Fig. 6A–C), all 84 neurons are included for a total of 269,687 fixations. For first fixations to an option that would be chosen (Fig. 6C), 55 neurons were included (accounting for 29,162 of 41,521 such fixations across all neurons). For first fixations to an option that would not be chosen (Fig. 6C), 60 neurons were included (accounting for 16,976 of 23,413 such fixations across all neurons). For the first fixation to the first-fixated option (Extended Data Fig. 3A), 49 neurons are included (39,640 of 64,934 such fixations). For the second fixation to the first-fixated option (Extended Data Fig. 3B), 77 neurons are included (49,616 of 54,345 such fixations). For the first fixation to the second-fixated option (Extended Data Fig. 3C), 76 neurons are included (51,883 of 57,288 such fixations). For the second fixation to the second-fixated option (Extended Data Fig. 3D), 64 neurons are included (30,694 of 40,966 such fixations).

### 4.10 Classification Based on Demixed PCs

Each attention, decision, value, and interaction dPC was used as a linear classifier for the corresponding dPC factor, following the procedure of Kobak et al. [23]. That is, we used 100 iterations of stratified MonteCarlo leave-group-out cross-validation. For a given dPC, the mean value of the dPC at each time point is computed for each condition. Test trials (left-out from the mean computation) are then projected onto the dPC and classified at each time point according to the closest condition mean. This gives the time intervals during which the respective factor can be reliably extracted from single-trial activity. Those time intervals are shown as color-coded horizontal bars at the bottom of the panels showing the time-dependent dPC components in Figs 4 and 6 and Extended Data Fig. 3.

### 4.11 Permutation Tests of Demixed PCs

A permutation test was used to assess significant differences between explained variance assigned to task factors using dPCA on different subsets of the data (i.e. comparing trials with four or fewer fixations to trials with five or more fixations as shown in Fig. 4D,E and comparing the different fixations to attributes of options shown in Extended Data Fig. 3). All data for each task condition to be compared were combined into one larger array (3 relative value bins-by-2 directions (chosen)-by-850 ms-by-trial number for the combined data set array for the response-aligned analysis in Fig. 4 and 4 relative value bin-by- 2 directions (attended)-by-250 ms-by-fixation number for the fixation-aligned analysis in Extended Data Fig. 3). These combined arrays were then randomly split 1000 times and subjected to dPCA to obtain a distribution of the difference in total explained variance assigned to each factor between the two split data sets. The true difference between the task conditions of interest for each parameter was then compared to this distribution. Differences were designated as significant if the absolute value of the difference for a given pair of task conditions was greater than 95% of the differences between random splits of the combined array (*p <* 0.05).

The same permutation testing procedure was also used to assess significant differences between the top-ranked decision-related dPC in trials with for or fewer fixations and trials with five or more fixations (see the 2.2 section). Four metrics were computed based on these components, after averaging out the value factor: 1) the maximum separation at any timepoint between trials when the PD option was chosen and trials when the NPD option was chosen, 2) the time at which this maximum separation occurred, 3) the slope of a line fit to the component between a prior timepoint and the point of maximum separation for trials when the PD option was chosen, and 4) the slope of a line fit to the component between a prior timepoint and the point of maximum separation for trials when the NPD option was chosen. For the response-aligned analysis, this prior timepoint was chosen to be 550 ms before the response. For the first fixation-aligned analysis, this prior timepoint was chosen to be the beginning of the first fixation. Differences in these metrics between the short and long trials were designated as significant if the difference was greater than 95% of the differences between random splits of the combined array (*p <* 0.05).

## Extended Data

**Table EDT.1:**
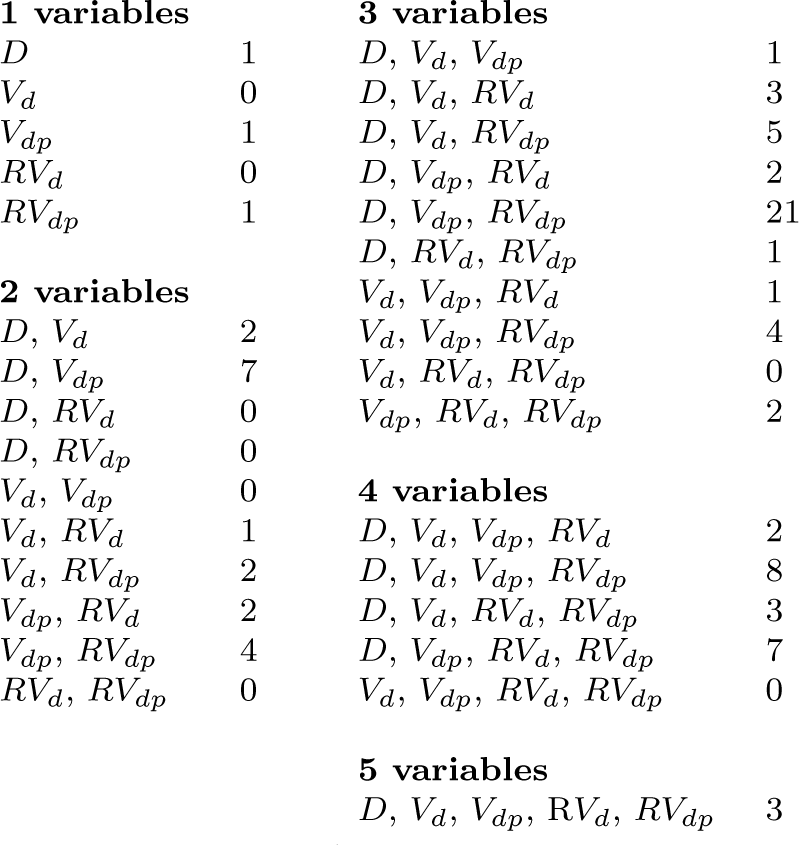
Number of preSMA neurons encoding the chosen option and different forms of the value of the chosen option at the time of the response in best-fit GLM.

|  |  |  |  |
| --- | --- | --- | --- |
| **1 variables** |  | **3 variables** |  |
| $D$ | 1 | $D, V_d, V_{dp}$ | 1 |
| $V_d$ | 0 | $D, V_d, RV_d$ | 3 |
| $V_{dp}$ | 1 | $D, V_d, RV_{dp}$ | 5 |
| $RV_d$ | 0 | $D, V_{dp}, RV_d$ | 2 |
| $RV_{dp}$ | 1 | $D, V_{dp}, RV_{dp}$ | 21 |
| **2 variables** | | $D, RV_d, RV_{dp}$ | 1 |
| $D, V_d$ | 2 | $V_d, V_{dp}, RV_d$ | 1 |
| $D, V_{dp}$ | 7 | $V_d, V_{dp}, RV_{dp}$ | 4 |
| $D, RV_d$ | 0 | $V_d, RV_d, RV_{dp}$ | 0 |
| $D, RV_{dp}$ | 0 | $V_{dp}, RV_d, RV_{dp}$ | 2 |
| $V_d, V_{dp}$ | 0 | **4 variables** | |
| $V_d, RV_d$ | 1 | $D, V_d, V_{dp}, RV_d$ | 2 |
| $V_d, RV_{dp}$ | 2 | $D, V_d, V_{dp}, RV_{dp}$ | 8 |
| $V_{dp}, RV_d$ | 2 | $D, V_d, RV_d, RV_{dp}$ | 3 |
| $V_{dp}, RV_{dp}$ | 4 | $D, V_{dp}, RV_d, RV_{dp}$ | 7 |
| $RV_d, RV_{dp}$ | 0 | $V_d, V_{dp}, RV_d, RV_{dp}$ | 0 |
|  |  | **5 variables** |  |
| | | $D, V_d, V_{dp}, RV_d, RV_{dp}$ | 3 |

**Fig. ED.1:**
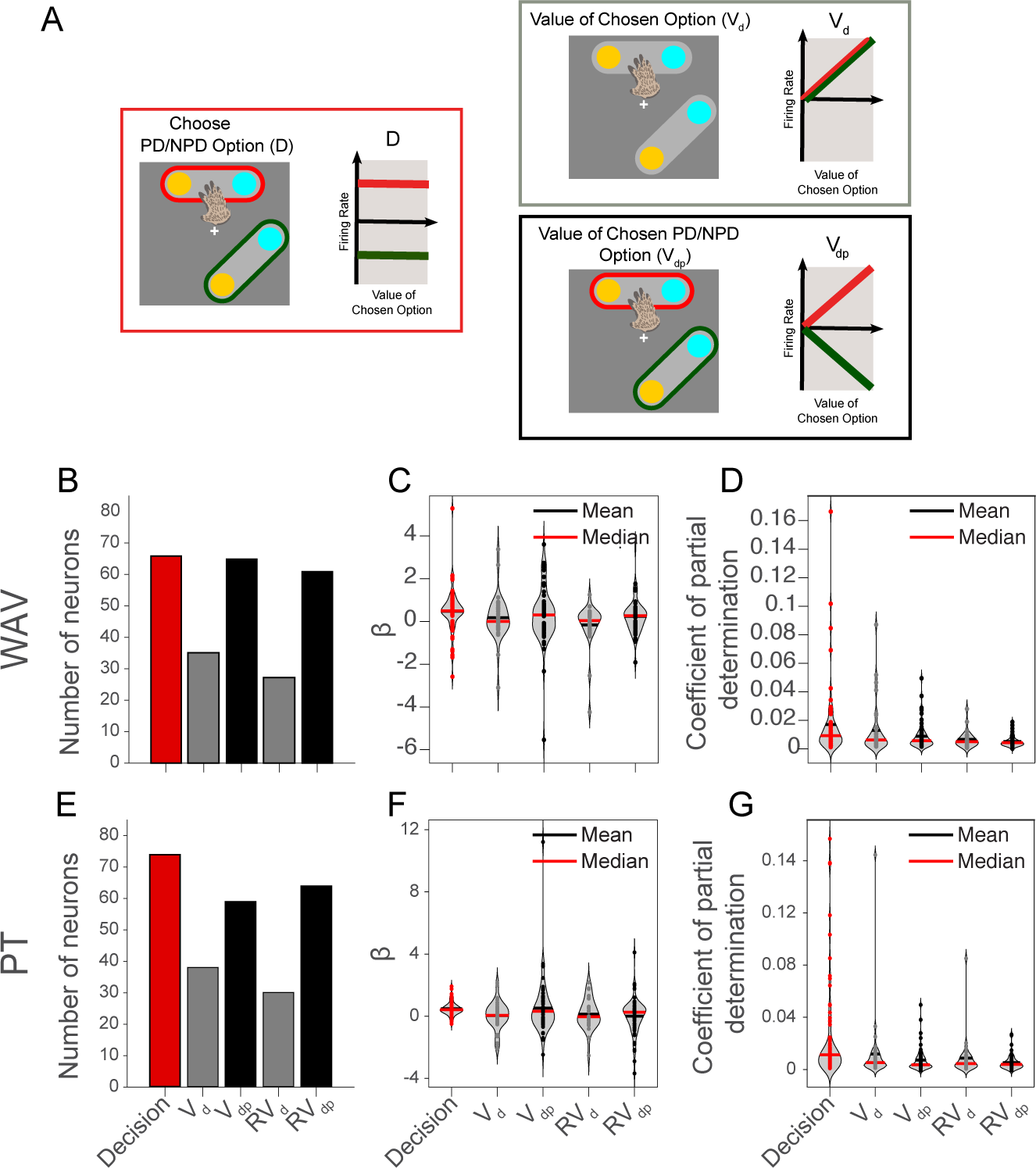
GLM analysis for response-aligned neural activity. (A) We analyzed the activity of preSMA neurons in the Response Period (200 ms before the beginning of arm movements ending in the selection of an option) using GLMs consisting of a range of different functional signals. We tested signals reflecting the decision (*D*, whether the preferred option is chosen) and a range of value signals. These value signals can reflect either the value of the chosen option (*V*) or the value difference between the chosen and non-chosen options (*RV*). For both value types, we tested both position-independent forms (*V_d_* and *RV_d_*) and forms that depend on whether the PD or NPD option was chosen (*V_dp_* and *RV_dp_*), see the Generalized Linear Model section in Methods for details. Each box contains a schematic depicting the dependency of each signal on the position of each option (indicated by the color border; red: PD, green: NPD) and whether or not it is chosen (indicated by the hand). To the right is a schematic of the expected firing rate as a function of the value of the chosen option (red: PD, green: NPD). Decision (D) indicates the spatial categorization of chosen options. The signal is binary, since attention is in either the PD or NPD. Value signals differ with respect to the framework for identifying the object they refer to. [Fig. ED.1 caption continues on next page.] Decision-dependent value signals (*V_d_* and *RV_d_*; gray box borders) indicate the value of options based on whether they are chosen or not (hand), independent of their position (no color borders for option symbols). Activity is expected to reflect value independent of whether the PD or NPD option is chosen. Decision and position-dependent value signals (*V_dp_* and *RV_dp_*; black box borders) indicate the value of options based on whether a specific position (PD/NPD; color borders) is chosen or not (hand). (B) The number of neurons whose best-fitting GLM includes specific predictor variable(s): Decision, *V_d_*, *V_dp_*, *RV_d_*, *RV_dp_*. The best model for each neuron was determined by comparing the AICs of models. The colors refer to the scheme used in Fig. ED.1A. All preSMA neurons encoded at least one of these variables and a large majority (81/84, 96%) encoded multiple variables. Decision (66/84, 79%), position-independent (*V_d_*or *RV_d_*, 49/84, 58%), and position-dependent value signals (*V_dp_* or *RV_d_p*, 77/84, 92%) were all strongly represented in preSMA population activity. (C) Coefficients *β* for each variable in the best-fit GLM for each neuron. (D) Coefficients of partial determination for each variable for all neurons. (E–G) GLM analysis for response-aligned neural activity using PT value instead of WAV for the value of an individual option. For a schematic representation of the decision- and position-dependent forms of value, see Fig. ED.1A. (E) The number of neurons whose best-fitting GLM includes specific predictor variable(s): Decision, *V_d_*, *V_dp_*, *RV_d_*, *RV_dp_*. The best model for each neuron was determined by comparing the AICs of models. The colors refer to the scheme used in Fig. ED.1A. (F) Coefficients *β* for each variable in the best-fit GLM for each neuron. (G) Coefficients of partial determination for each variable for all neurons.

**Fig. ED.2:**
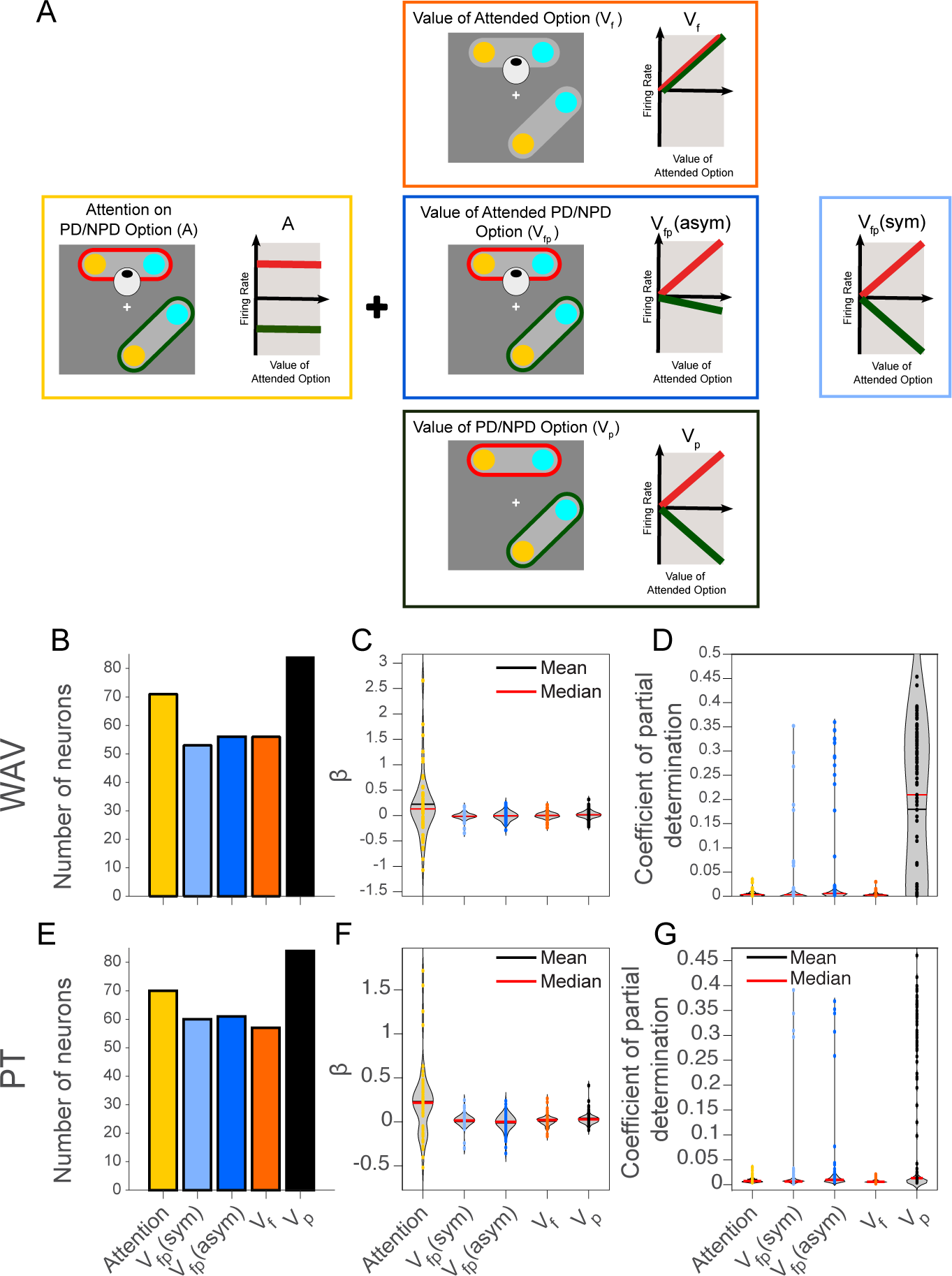
GLM analysis for fixation-aligned neural activity using PT value instead of WAV for the value of an individual option. For a schematic representation of the fixation- and position-dependent forms of value, see Fig. 5A. (E) The number of neurons whose best-fitting GLM includes specific predictor variable(s): Attention, *V_fp_*(symmetric), *V_fp_*(asymmetric), Vf, *V_p_*. The best model for each neuron was determined by comparing the AICs of models. The colors refer to the scheme used in Fig. 5A. (F) Coefficients *β* for each variable in the best-fit GLM for each neuron. (G) Coefficients of partial determination for each variable for all neurons.

**Fig. ED.3:**
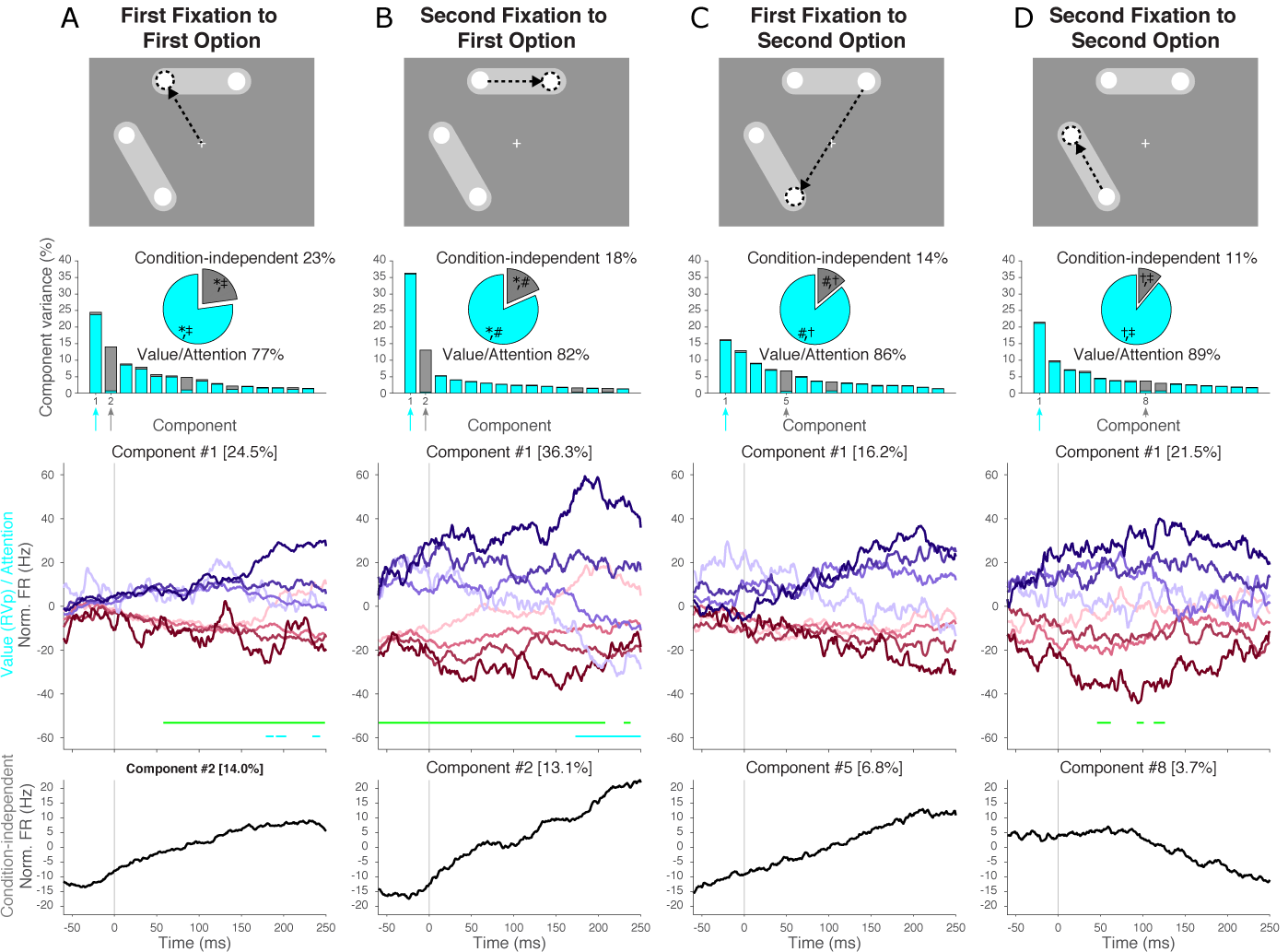
dPCA of Neural Activity during the Attribute Search Period across Fixation Sequence. Monkeys most commonly sampled the attributes from the two options by first investigating the attributes of one option before investigating the attributes of the other option (Fig. 2H). Shown are results from the dPCA analysis in Fig. 6A-C but separately for each of these four fixations. Each column shows, top to bottom: a schema of the fixation sequence, the first 15 demixed components and a pie chart showing the overall distribution of explained variance, and the projection of fixation-aligned neuronal spike densities onto the strongest functional components (indicated by the arrows; color code as in Fig. 6). (A) The first fixation to the first-fixated option. (B) The first fixation to the second-fixated attribute of the first-fixated option. (C) The first fixation to the second-fixated option. (D) The first fixation to the second-fixated attribute of the second-fixated option. In the pie charts, significant differences in total explained variance per factor between fixations are denoted by *∗* (first fixation to first option *vs.*second fixation to first option), # (first fixation to first option *vs.*second fixation to second option), *†* (first fixation to second option and second fixation to second option), and *‡* (first fixation to first option and second fixation to second option). Both factors were compared using a permutation test (see the Permutation Tests of Demixed PCs section in Methods) for panel A *vs.*B, B *vs.*C, C *vs.*D, and A *vs.*D. In all component 1 plots in (A–D), time periods of significant classification (see the Classification Based on Demixed PCs section in Methods) of the *RV_p_/A* factor are marked with a horizontal line near the bottom of the plot, see legend of Fig. 6 for color coding. Where no horizontal line is shown, the factor could not be significantly classified based the plotted component.

**Fig. ED.4:**
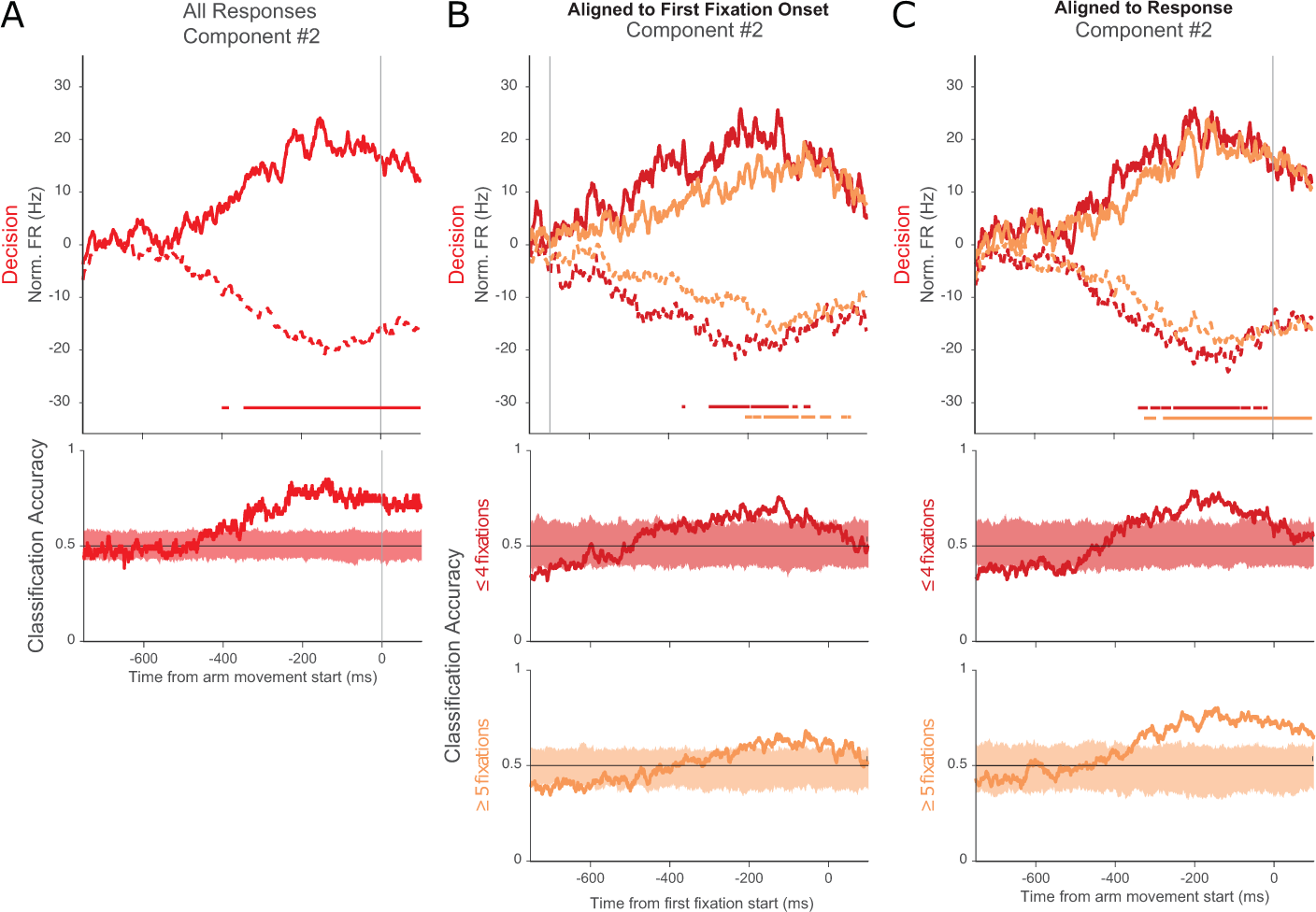
Classification accuracy for using de-mixed PCs as linear decoders of the decision from neural activity during the response period. Each column shows, at top: the projection of neuronal spike densities onto the strongest functional components with time periods of significant classification (see section 4.9) of the decision factor marked with a horizontal line near the bottom of the plot, and at bottom, the classification accuracy at each time point superimposed on a shaded region corresponding to 95% of classification accuracy for randomly shuffled trials. (A) All trials, aligned to response. (B) Separately for trials with four or fewer (dark red) and trials with five or more (light red) fixations, aligned the the beginning of the first fixation. (C) Separately for trials with four or fewer (dark red) and trials with five or more (light red) fixations, aligned the the beginning of the response.

**Fig. ED.5:**
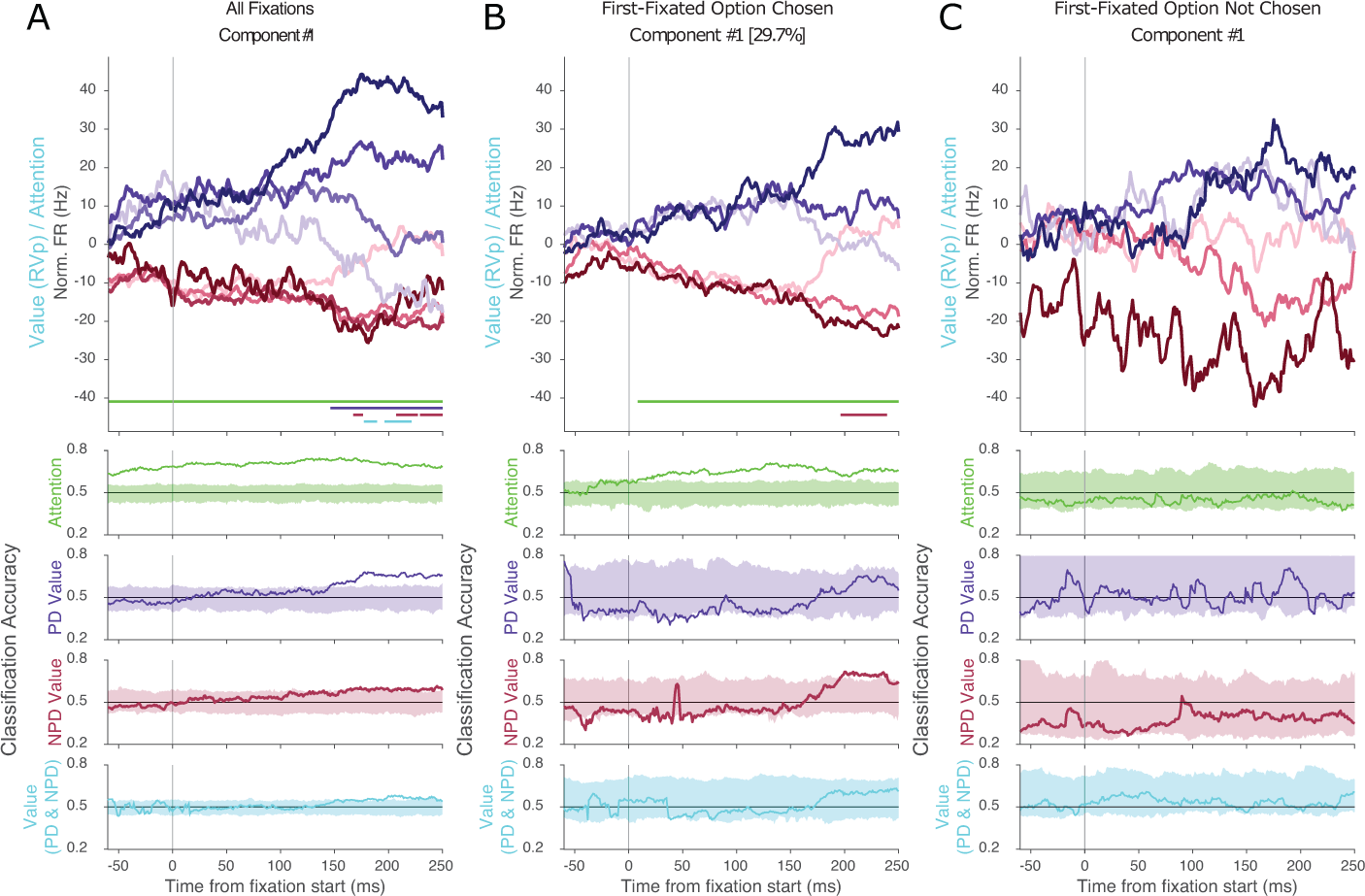
Classification accuracy for using de-mixed PCs as linear decoders of neural activity during the attribute search period WAV for option value. Each column shows, at top: the projection of fixation-aligned neuronal spike densities onto the strongest functional components with time periods of significant classification (see section 4.9) of the *RV_p_/A* factor marked with a horizontal line near the bottom of the plot (color code as in Fig. 6), and at bottom, the classification accuracy at each time point superimposed on a shaded region corresponding to 95% of classification accuracy for randomly shuffled trials for each of four binary classifications: attention (PD *vs.*NPD), PD value (high *vs.*low), NPD value (high *vs.*low), and value of the fixation option regardless of position (high *vs.*low). (A) All fixations. (B) Fixations to the first option when that option would be chosen. (C) Fixations to the first option when that option would not be chosen In all component 1 plots in (A–C), time periods of significant classification (see section 4.9) of the *RV_p_/A* factor are marked with a horizontal line near the bottom of the plot, see legend of Fig. 6 for color coding. Where no horizontal line is shown, the factor could not be significantly classified based the plotted component.

**Fig. ED.6:**
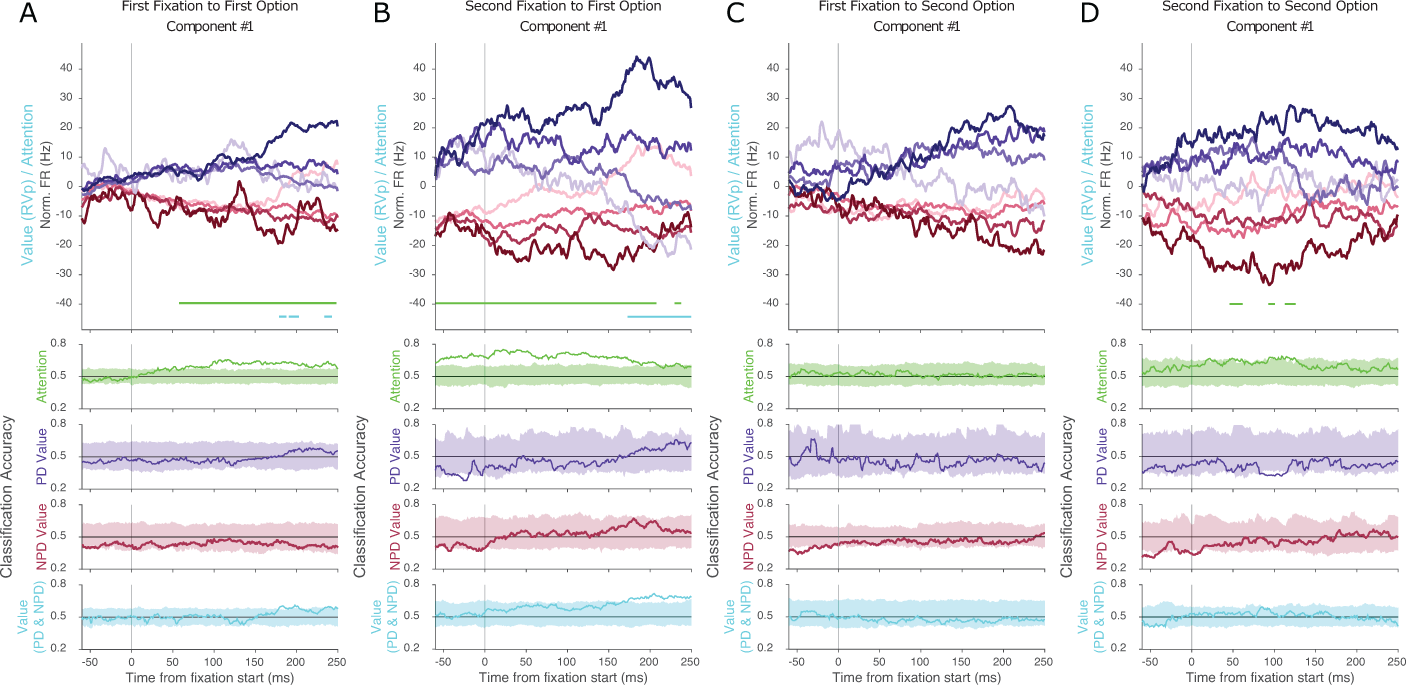
Classification accuracy for using de-mixed PCs as linear decoders of neural activity during the attribute search period WAV for option value, separated by specific fixations to attributes of options. Each column shows, at top: the projection of fixation-aligned neuronal spike densities onto the strongest functional components with time periods of significant classification (see section 4.9) of the *RV_p_/A* factor marked with a horizontal line near the bottom of the plot (as in Fig. 7; color code as in Fig. 6), and at bottom, the classification accuracy at each time point superimposed on a shaded region corresponding to 95% of classification accuracy for randomly shuffled trials for each of four binary classifications: attention (PD *vs.*NPD), PD value (high *vs.*low), NPD value (high vs. low), and value of the fixation option regardless of position (high vs. low). (A) The first fixation to the first-fixated option. (B) The first fixation to the second-fixated attribute of the first-fixated option. (B) The first fixation to the second-fixated option. (D) The first fixation to the second-fixated attribute of the second-fixated option. In all component 1 plots in (A–D), time periods of significant classification (see section 4.9) of the *RV_p_/A* factor are marked with a horizontal line near the bottom of the plot, see legend of Fig 6 for color coding. Where no horizontal line is shown, the factor could not be significantly classified based the plotted component.

**Fig. ED.7:**
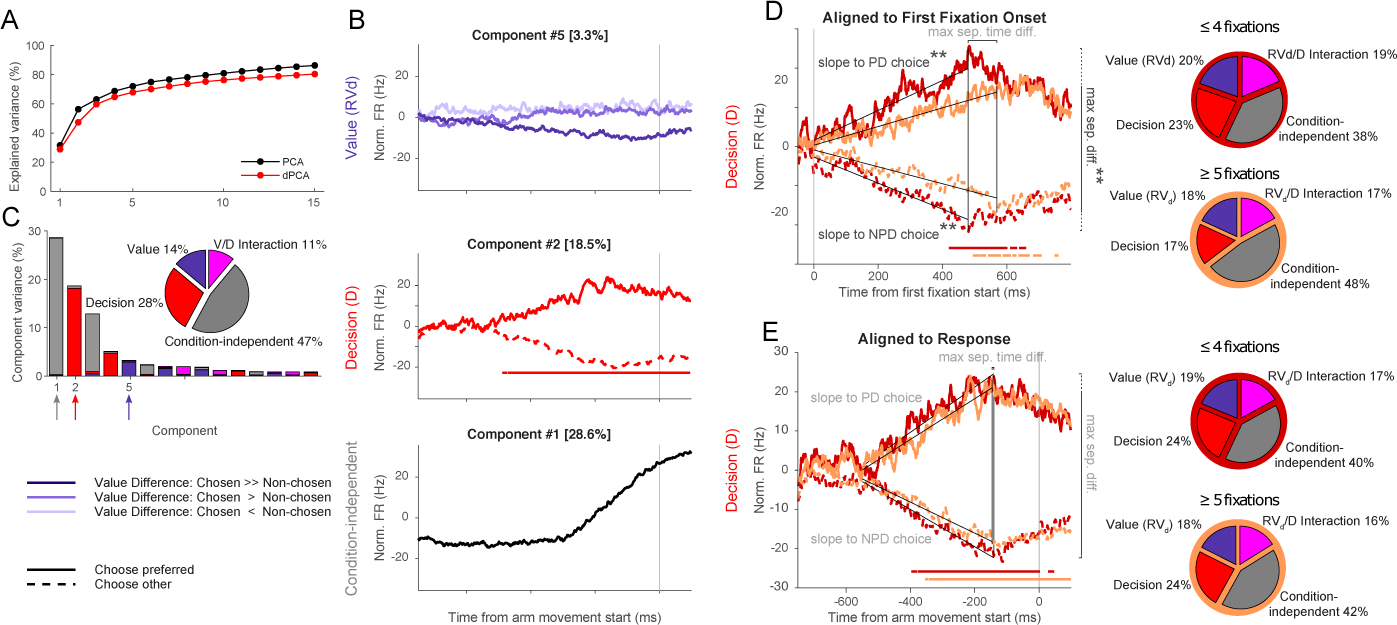
dPCA of Neural Activity during the Response Period using PT value instead of WAV for the value of an individual option. Here, dPCA was used to identify the components related to two key factors: value (*RV d*, the difference between the subjective values of the chosen and non-chosen options) and decision (the position, preferred or non-preferred, of the chosen option), as well as the interaction of these factors and condition-independent components. (A) Explained variance across components for dPCA and PCA. (B) Histogram of the first 15 components with regard to the value and decision factors, their interaction, and condition-independent variance. The pie chart shows the overall distribution of explained variance across these factors. Components 1, 2, and 5 (arrows under abscissa; colors correspond to those in pie chart and in panel C) are the largest condition-independent, decision-related, and value-related components, respectively. (C) Time course of the top-ranked value-related, decision-related, and condition-independent components, i.e. the projections of fixation-aligned neuronal spike densities onto the individual components. For Components 2 and 6, periods of significant classification (see section 4.9) are denoted with a black horizontal line at the bottom of each plot. (D–E) Top-ranked decision components for dPCA on trials with fewer (*≤* 4; dark red lines) and more (*≤* 5; light red lines) fixations. (D) dPCA for data aligned to first fixation onset. (E) dPCA for data aligned to arm movement. In D and E, periods of significant classification of the decision factor based on the plotted component are shown as horizontal lines in the same color as the corresponding time course plot.

**Fig. ED.8:**
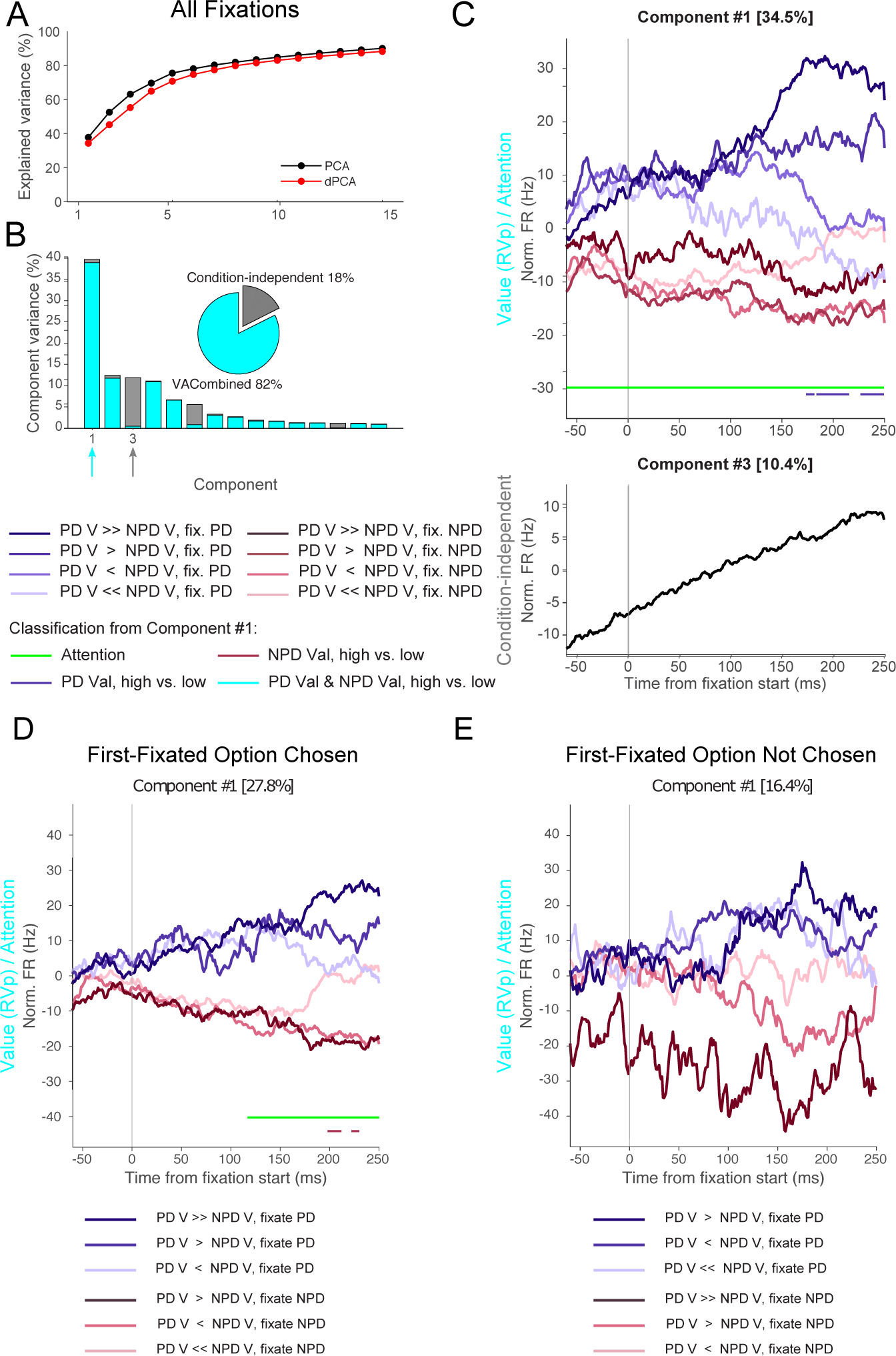
dPCA of Neural Activity during the Attribute Search Period using PT value instead of WAV for the value of an individual option. As in Fig. 6, we demix activity using only one factor, called *RV_p_/A* (plus the condition-independent factor) which consists of the relative value *RV_p_* multiplexed with position-defined relative value *A*. This factor has eight states, see legend under panel B. (A) Explained variance across components for dPCA and PCA. [Fig. ED.8 caption continues on next page.] (B) Histogram of the first 15 components with regard to the *RV_p_/A* factor and condition-independent variance. The pie chart shows the overall distribution of explained variance across these factors. Components 1 and 3 (arrows under abscissa; colors correspond to those in pie chart and in panels C-E) are the largest components of the *RV_p_/A* factor and the condition-independent activity, respectively. (C) Upper panel: Time course of the eight states of *RV_p_/A*, i.e. the projections of fixation-aligned neuronal spike densities onto the individual components. For Component 1, periods of significant classification (see section 4.9) are denoted with a horizontal line at the bottom of each plot, see legend for color coding. Where no horizontal line is shown, the state could not be significantly classified based on component 1. Lower panel: Time course of projection onto the condition-independent factor. (D,E) Same as in A but only for first fixations in each trial. Here reduced sets of states for the *RV_p_/A* factor are used, see legends under panels D and E. (D) Trials in which the first fixated option was chosen. (E) Trials in which the first fixated option was not chosen.

**Fig. ED.9:**
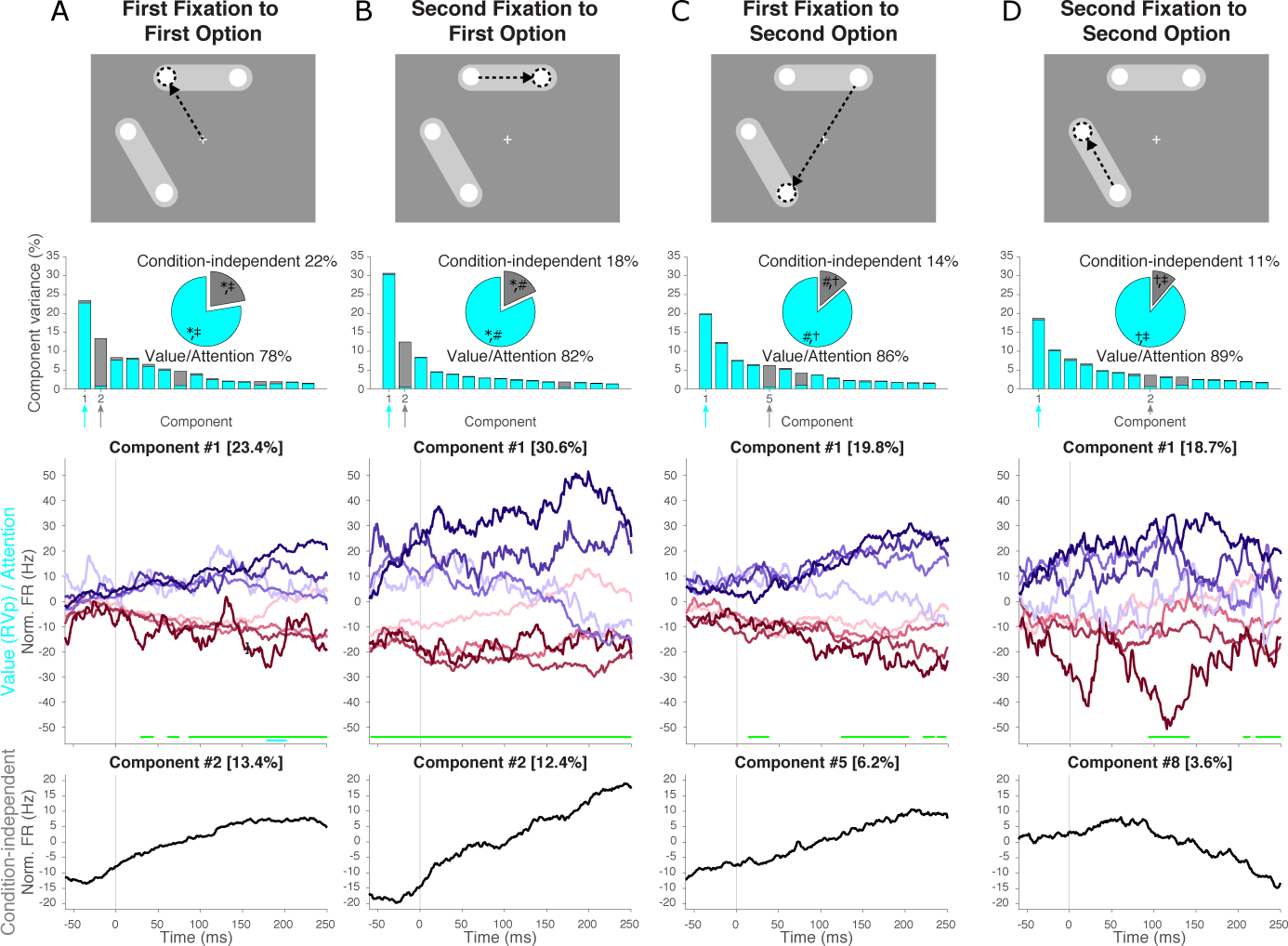
dPCA of neural activity during attribute search period using PT instead of WAV for option value, separated by specific fixations to attributes of options. As in Fig. 7, neural activity is analyzed separately for each of the four fixations to a new attribute. Each column shows, top to bottom: a schema of the fixation sequence, the first 15 demixed components and a pie chart showing the overall distribution of explained variance, and the projection of fixation-aligned neuronal spike densities onto the strongest functional components (indicated by the arrows; color code as in Fig.6). (A) The first fixation to the first-fixated option. (B) The first fixation to the second-fixated attribute of the first-fixated option. (C) The first fixation to the second-fixated option. (D) The first fixation to the second-fixated attribute of the second-fixated option. In all component 1 plots in (A–D), time periods of significant classification (see section 4.9) of the *RV_p_/A* factor are marked with a horizontal line near the bottom of the plot, see legend of Fig 6 for color coding. Where no horizontal line is shown, the factor could not be significantly classified based the plotted component.

## Supplementary Information

### 1 Behavioral Modeling of Risky Choices

The monkeys needed to integrate the amount and probability information to assign a subjective value to the option. We considered four possible models of this process (for details see Methods). In two models, the option-value was a multiplicative function of the attribute-values. The expected value (EV) model assumes that monkeys use linear representations of reward amount and probability. In contrast, the prospect theory (PT) model assumes monkeys use nonlinear functions that transform objective amount and probability to their corresponding subjective mappings (i.e. utility *u*(*a*) and weighted probability *w*(*p*) [1, 2]). In two other models, the option-value was an additive function of the attribute-values. The additive value (AV) model assumes monkeys calculated the value of an option by summing the normalized magnitudes of its corresponding amount and probability attributes. The weighted additive value (WAV) model extends the AV model by allowing that the two attributes have different significance for the monkeys. Thus, we apply weights to one of the attributes to obtain a weighted additive value of the option. We compared the behavioral fits using prediction accuracy and the Bayesian Information Criterion (BIC).

**Table ST.1.** Choice prediction accuracy for all trials. Lower BIC values generally indicate better models, but are not comparable across monkeys due to different trial counts.

| model type | # free param. | monkey H prediction accuracy | monkey I prediction accuracy | monkey H BIC | monkey I BIC |
| --- | --- | --- | --- | --- | --- |
| EV | 0 | 81.99% | 78.78% | 8482 | 4118 |
| PT | 2 | 84.96% | 81.74% | 7622 | 3886 |
| AV | 0 | 80.04% | 80.19% | 8915 | 3819 |
| WAV | 1 | 84.83% | 83.04% | 7506 | 3346 |

Models with different weights of attributes *a* and *p* (PT and WAV) outperformed their corresponding models without this assumption (EV and AV; Table ST.1). Monkeys were generally risk-seeking, corresponding to a higher weighting of amount relative to probability in both PT (in Eq. 2: *α* = 1.46 and 1.7 for monkey I and H, respectively; in Eq. 3: *γ* = 0.79 and 0.88 for monkey I and H, respectively) and WAV (*w_a_* = 1.86 and 1.99 for monkeys I and H, respectively) consistent with past work ([2–5]).

It is less clear whether an additive or a multiplicative rule better captures the monkeys’ preferences. The best additive model (WAV) and the best multiplicative model (PT) predicted choice nearly equally well (Table 1). Additive and multiplicative value estimates were strongly correlated (corr. PT vs WAV: *R*^2^ = 0.66 for monkey H; *R*^2^ = 0.68 for monkey I). We therefore analyzed behavior specifically on trials in which the PT and the WAV model made different predictions about which option should be preferred. Again, the results did not strongly support one model over the other (correctly predicted by PT: 51.04% for monkey H; 43.6% for monkey I; the remainder was better predicted by WAV). We decided to use the WAV model over the PT model to estimate subjective value in the neuronal analysis because of its better fit as measured by BIC. However, we also repeated all analyses using value estimates derived from the PT model and found matching results (Extended Data).

### 2 Behavioral Evidence that monkeys understood meaning of symbolic cues indicating attribute magnitude

The monkeys understood the meaning of the symbolic cues. When both options were rewarded with equal probability, they chose more often the option with a higher reward amount (for both monkeys, P(choose more valuable option) = 0.9 *±* 0.01)). Likewise, when both options offered equal reward amounts, monkeys were more likely to choose the option with higher reward probability (Monkey I: P(choose more valuable option) = 0.75 *±* 0.01; Monkey H: P = 0.77 *±* 0.02). These results indicate that the monkeys correctly associated the two colored circles as representing two different attributes of one object and preferred symbols corresponding to higher-valued attributes over lower-valued ones.

**Fig. S.1.**
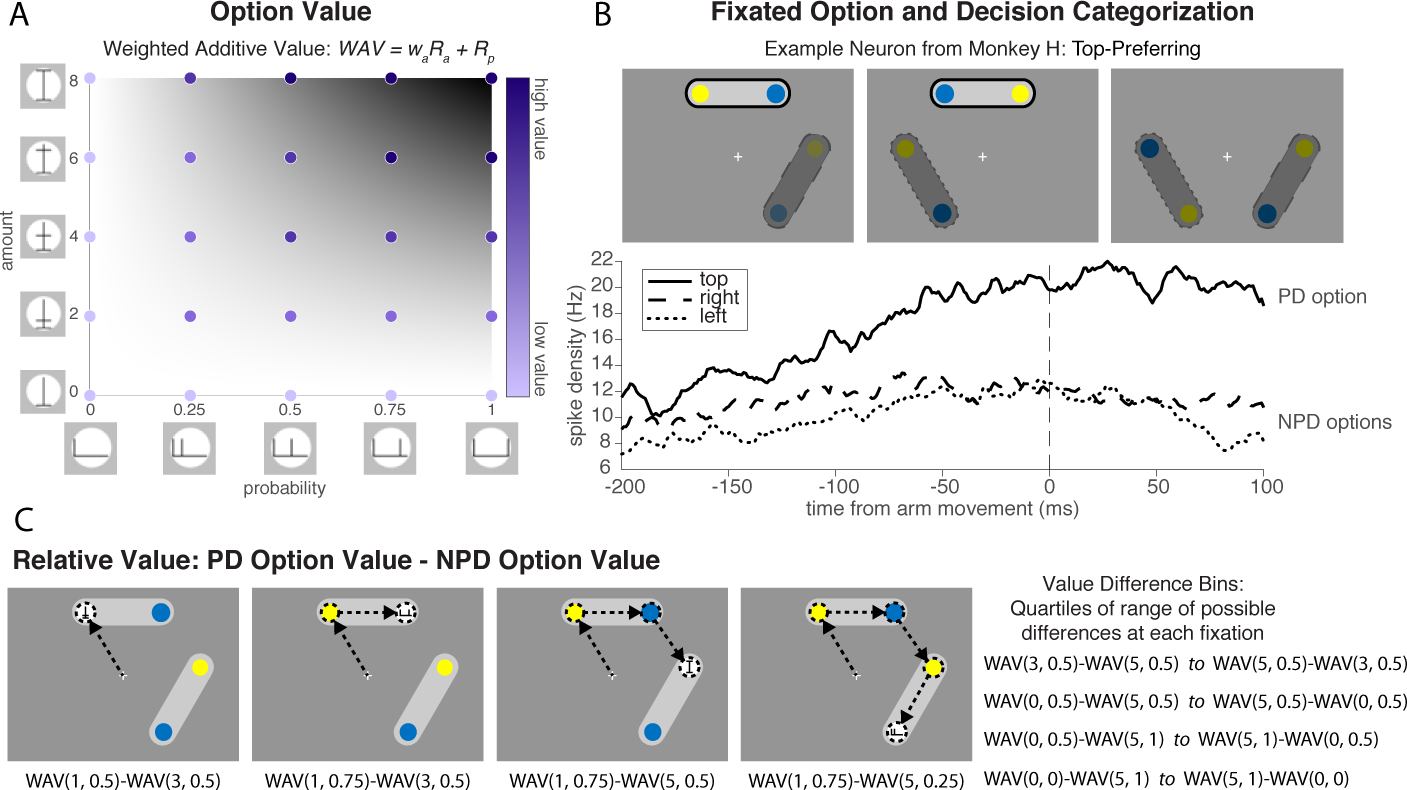
Choice-related variables in preSMA neurons. (A) Value space for monkey H estimated by a weighted additive value model (the space for monkey I is equivalent but only has four values along each axis). All values were sorted into four value bins. (B) To categorize *Attention* and *Decision* signals, the preferred direction for each neuron was first found based on the mean activity of each neuron in the 200 ms period prior to the arm movement response. In this example, neurons whose neuronal activities were significantly higher when the joystick was moved to the top than to the right or left during the arm movement period, ‘top’ is defined as its preferred direction (PD) while ‘right’ and ‘left’ are defined as its non-preferred direction (NPD). The *Attention* signal was assigned the value 1 when the current fixation was to the top option (PD; bright bar with solid-line boundary) and -1 when the current fixation was to the right or left option (NPD; muted-color bars with dashed-line boundaries). The *Decision* signal was assigned 1 when the joystick was moved to the top option (PD; solid-line) and -1 when the joystick was moved to the right or left (NPD; dashed-lines). (C) The *V alue* signal was computed at each fixation according to the information that had been collected by the monkey up to that point.

